# Observation of coordinated cotranscriptional RNA folding events

**DOI:** 10.1101/2023.02.21.529405

**Authors:** Courtney E. Szyjka, Eric J. Strobel

## Abstract

RNA begins to fold as it is transcribed by an RNA polymerase. Consequently, RNA folding is constrained by the direction and rate of transcription. Understanding how RNA folds into secondary and tertiary structures therefore requires methods for determining the structure of cotranscriptional folding intermediates. Cotranscriptional RNA chemical probing methods accomplish this by systematically probing the structure of nascent RNA that is displayed from RNA polymerase. Here, we have developed a concise, high-resolution cotranscriptional RNA chemical probing procedure called <u>T</u>ranscription <u>E</u>longation <u>C</u>omplex RNA structure <u>prob</u>ing-<u>M</u>ultilength (TECprobe-ML). We validated TECprobe-ML by replicating and extending previous analyses of ZTP and fluoride riboswitch folding, and mapped the folding pathway of a ppGpp-sensing riboswitch. In each system, TECprobe-ML identified coordinated cotranscriptional folding events that mediate transcription antitermination. Our findings establish TECprobe-ML as an accessible method for mapping cotranscriptional RNA folding pathways.

## Introduction

RNA begins to fold as it emerges from an RNA polymerase (RNAP) during transcription^1, 2^. Consequently, the rate and 5’ to 3’ direction of transcription directly influence RNA folding, and nascent RNA structures can interact with RNAP to control transcription^3–5^. Although the role of transcription in RNA structure formation and the coordination of cotranscriptional processes has been established for decades^3, 6, 7^, our ability to measure cotranscriptional RNA folding remained limited until recently. In the past several years, the development of methods that can monitor cotranscriptional RNA structure formation with high temporal and spatial resolution has rapidly advanced our ability to characterize cotranscriptional RNA folding mechanisms. These complementary approaches include applications of single-molecule force spectroscopy^8, 9^ and single-molecule FRET^10–13^, which measure cotranscriptional RNA folding with high temporal resolution, and the application of high-throughput RNA chemical probing to map the structure of cotranscriptionally folded intermediate transcripts at nucleotide resolution^14–16^.

High-throughput RNA chemical probing methods characterize the structure of complex RNA mixtures at nucleotide resolution and are compatible with diverse experimental conditions^17–19^. This experimental flexibility enabled the development of RNA chemical probing assays in which *E. coli* RNAP is distributed across template DNA so that cotranscriptionally folded intermediate transcripts can be chemically probed^14–16^. This approach systematically maps the structure of potential cotranscriptional RNA folding intermediates in the context of RNAP and identifies when secondary and tertiary structures can fold within an RNA folding pathway. Despite their utility for mapping RNA folding pathways, cotranscriptional RNA chemical probing experiments can be challenging to execute because existing protocols are relatively complex.

We have developed a concise, SHAPE-MaP-based^20^ (<u>s</u>elective 2’-<u>h</u>ydroxyl <u>a</u>cylation analyzed by <u>p</u>rimer <u>e</u>xtension and <u>m</u>utational <u>p</u>rofiling) procedure for multi-length cotranscriptional RNA chemical probing assays called TECprobe-ML (<u>T</u>ranscription <u>E</u>longation <u>C</u>omplex RNA structure <u>prob</u>ing-<u>M</u>ultilength). TECprobe-ML was designed with the specific objectives of standardizing cotranscriptional RNA structure probing assays and eliminating target-dependent variability in experiment outcomes. At the experimental level, this is accomplished through a streamlined procedure that yields sequencing libraries with a ∼90% alignment rate and guarantees even sequencing coverage across the length of each intermediate transcript. At the computational level, we implemented a data smoothing procedure to overcome experimentally irresolvable biases in the sequencing coverage of intermediate transcripts. We validated TECprobe-ML by repeating and extending previous analyses of the *Clostridium beijerinckii* (*C. beijerinckii*, *Cbe*) *pfl* ZTP^21^ and *Bacillus cereus* (*B. cereus*, *Bce*) *crcB*^22^ riboswitches. In addition to replicating all prior observations, TECprobe-ML resolved biologically relevant folding events that were not previously detectable. We then used TECprobe-ML to map the folding pathway of a ppGpp-sensing riboswitch^23^ from *Clostridiales bacterium oral taxon* 876 str. F0540 (*C. bacterium*, *Cba*). Our analysis determined that (i) the *Cba* ppGpp riboswitch folds using a branched pathway in which mutually exclusive intermediate structures converge to form the ppGpp aptamer, and (ii) ppGpp binding coordinates extensive long-range contacts between ppGpp aptamer subdomains. Together, these advances establish TECprobe-ML as a high-performance and broadly accessible cotranscriptional RNA structure probing method.

## Results

### Optimization and benchmarking of TECprobe-ML

TECprobe-ML is designed to simplify and standardize cotranscriptional RNA structure probing experiments. This is primarily accomplished by including an established 5’ structure cassette^24^ (referred to here as SC1) upstream of the target RNA sequence and detecting chemical adducts by mutational profiling using the SHAPE-MaP^20^ strategy (Figure 1). Mutational profiling uses error-prone reverse transcription to encode the location of RNA chemical adducts, which increase the error rate of reverse transcriptase, as mutations in cDNA^20^. Because SHAPE-MaP can encode reactivity information in full-length cDNA molecules, the inclusion of a 5’ structure cassette upstream of the target RNA facilitates direct amplification of cDNA^20^. In contrast, the original Cotranscriptional SHAPE-Seq procedure required the ligation of an adapter to the cDNA 3’ end because chemical adducts were detected as truncated cDNA^14, 15^. This ligation is prone to sequence bias that distorts the reactivity measurement and can cause the formation of adapter dimer, which reduces the number of usable sequencing reads. The removal of adapter dimer by size selection also depletes short cDNA products, resulting in poor resolution of 3’-proximal RNA structures. Consequently, eliminating the need to ligate an adapter to the cDNA 3’ end resolves every limitation of the original Cotranscriptional SHAPE-Seq procedure that was caused by adapter dimer: TECprobe-ML libraries have a ∼90% alignment rate (Figure 2a), and resolve 3’-proximal structures that were not detectable previously (as is shown for the ZTP and fluoride riboswitches below).

**Figure 1.**
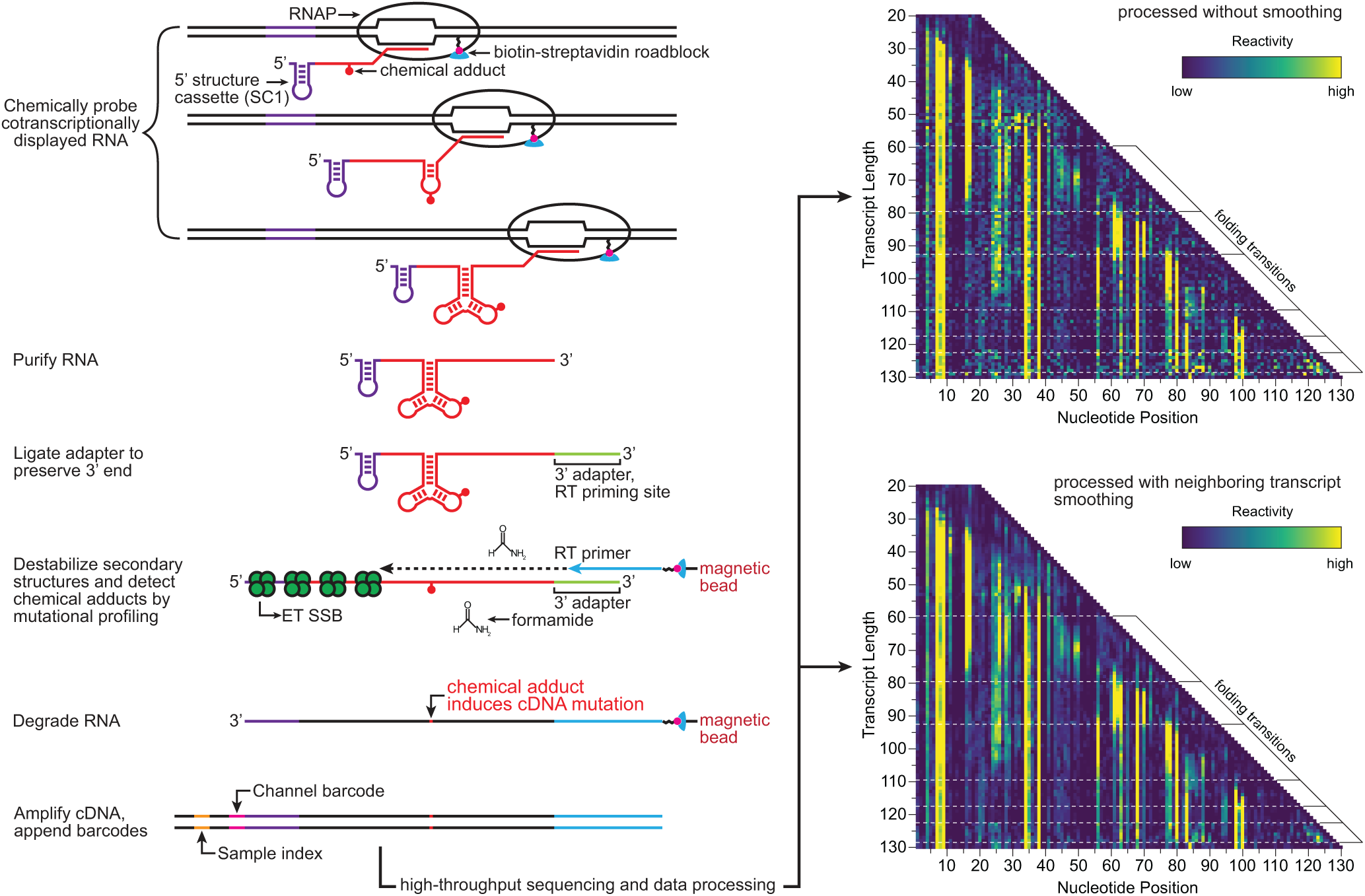
Overview of TECprobe-ML. Randomly biotinylated DNA templates are *in vitro* transcribed to generate cotranscriptionally folded intermediate transcripts displayed from *E. coli* RNAP. The cotranscriptionally displayed intermediate transcripts are chemically probed and purified. An adapter is ligated to the RNA 3’ end and used to prime solid-phase error-prone reverse transcription. Full-length cDNA is amplified using primers that anneal to the adapter and the 5’ structure cassette. Following high-throughput sequencing, data are processed using fastp^56^, TECtools, and ShapeMapper2^57^ either with or without neighboring transcript smoothing. SC1, structure cassette 1; RT, reverse transcription; ET SSB, extremely thermostable single-stranded binding protein.

**Figure 2.**
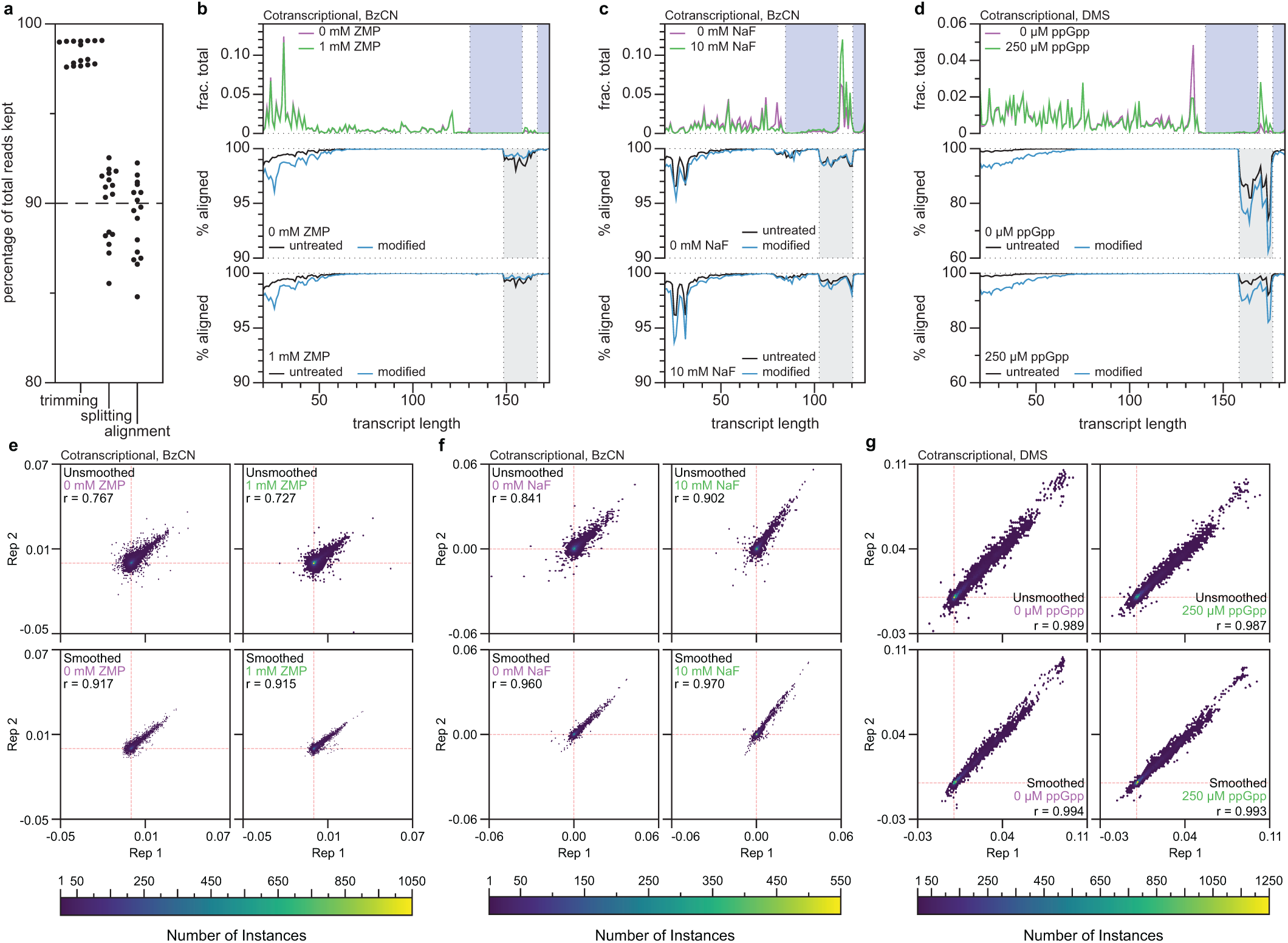
TECprobe-ML performance benchmarks. **a,** Percentage of total reads kept after adapter trimming, splitting fastq files by transcript length and channel barcode, and sequencing read alignment**. b-d,** Plots showing the fraction of aligned reads that mapped to each transcript length (top) and the percentage of split reads that aligned to each transcript length (middle, bottom) for the *Cbe pfl* ZTP riboswitch (BzCN) (**b**), the *Bce crcB* fluoride riboswitch (BzCN) (**c**), and the *Cba* ppGpp riboswitch (DMS) (**d**) datasets. The blue shading in the top plot indicates transcripts that were not enriched by biotin-streptavidin roadblocks. The grey shading in the middle and lower plots indicates transcripts in which alignment was lower due to the presence of a nucleic acid species that is kept during fastq splitting but does not align. **e-g**, Hexbin plots comparing the reactivity of replicates for the *Cbe pfl* ZTP (BzCN) (**e**), *Bce crcB* fluoride (BzCN) (**f**), and *Cba* ppGpp (DMS) (**g**) riboswitches for both unsmoothed (top) and smoothed (bottom) datasets. Data were plotted with a grid size of 75 by 75 hexagons, and the depth of overlapping data points are indicated by the heatmap. Red dashed lines indicate the position of 0 for each axis. BzCN, benzoyl cyanide; DMS, dimethyl sulfate.

In addition to eliminating the need for cDNA ligation, we optimized several steps of the TECprobe-ML library prep. First, ligation of an RNA 3’ adapter is performed using the reverse complement of the Illumina TruSeq Small RNA Kit RA5 adapter so that Read 1 begins with high-complexity sequence, which is necessary for cluster identification during Illumina sequencing. Second, after the RNA 3’ adapter ligation, RNA is purified using solid-phase reversible immobilization (SPRI) to remove most excess adapter prior to reverse transcription (Supplementary Figures 1 and 3a). Third, reverse transcription is performed as a solid-phase reaction to simplify cDNA library purification (Figure 1). Trace amounts of excess RNA 3’ adapter in the solid-phase reverse transcription reaction caused the formation of an RT primer product due to template switching (Supplementary Figures 2a, b and 3a). However, this product was not amplifiable and therefore had no effect on the procedure (Supplementary Figure 2c). Fourth, reverse transcription is performed in the presence of thermostable single-stranded DNA binding protein (ET SSB) and formamide to eliminate RT primer dimer and promote full-length cDNA synthesis (Figure 1 and Supplementary Figure 3b). In combination, structuring full-length cDNA as an amplicon and the protocol optimizations described above enabled us to reduce the scale of the TECprobe-ML *in vitro* transcription reaction by 10-fold relative to the original Cotranscriptional SHAPE-Seq protocol.

**Figure 3.**
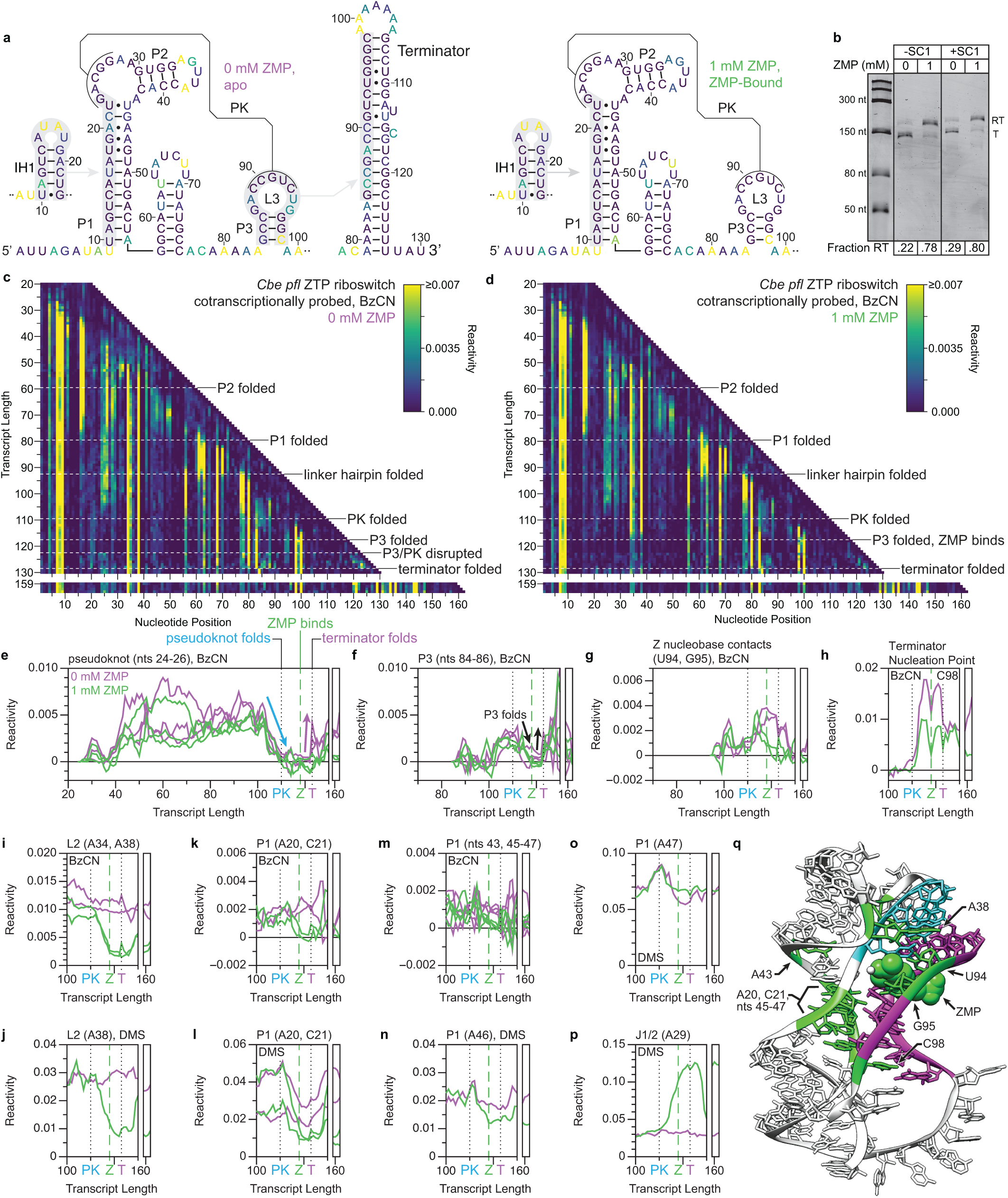
Cotranscriptional folding transitions of the *C. beijerinckii pfl* ZTP riboswitch. **a**, Secondary structures of *pfl* ZTP riboswitch folding intermediates colored by reactivity (IH1: BzCN, 41 nt; Apo/ZMP-bound aptamer: BzCN, 120 nt; terminator: DMS, 130 nt). **b**, Single-round transcription termination assays for the *pfl* ZTP riboswitch with and without SC1. Fraction readthrough is the average of two independent replicates. **c,d**, TECprobe-ML BzCN reactivity matrices for the *pfl* ZTP riboswitch with 0 mM and 1 mM ZMP. Transcripts 131- 158, which were not enriched, are excluded. Data are from two independent replicates that were concatenated and analyzed together. Reactivity is shown as background-subtracted mutation rate. **e-p**, Transcript length-dependent reactivity changes showing folding transitions. DMS data are from Supplementary Figure 7a, b. Black dotted lines indicate pseudoknot and terminator folding. Green dashed lines indicate ZMP binding. **q**, *Thermosinus carboxydivorans pfl* riboswitch structure (PDB 4ZNP)^27^. P3 (magenta), J1/2 PK (cyan), and ZMP- responsive nucleotides (green) are highlighted. BzCN, benzoyl cyanide; DMS, dimethyl sulfate; PK, pseudoknot; SC1, structure cassette 1; RT, readthrough; T, terminated.

The remaining limitation of cotranscriptional RNA structure probing procedures is that sequence and structural biases from the RNA 3’ adapter ligation can cause uneven representation of intermediate transcripts in the sequencing library^15^ (Figure 2b-d and Supplementary Figure 4a-c, upper plots). Because there is unlikely to be a general experimental solution to this limitation, we implemented a computational solution. Our approach is based on the observation that neighboring transcripts tend to have similar reactivity patterns and background mutation rates (Supplementary Figure 5). Therefore, concatenating sequencing reads from transcripts *n*-1, *n*, and *n*+1 in overlapping windows across a cotranscriptional RNA structure probing data set is expected to preserve transcript-length-dependent reactivity trends while boosting the effective sequencing depth for each length *n*. The use of neighboring transcript smoothing improved the correlation between replicate data sets, particularly when BzCN was used (Figure 2e-g and Supplementary Figure 4d-f). Comparisons of unsmoothed and smoothed data sets for each riboswitch RNA assessed in this work are shown in Supplementary Figure 6.

### Validation: The *C*. *beijerinckii pfl* ZTP riboswitch

Strobel et al. previously used Cotranscriptional SHAPE-Seq to map the folding pathway of the *Cbe pfl* ZTP riboswitch^25^, which controls gene expression in response to the purine metabolites ZMP and ZTP using a transcription antitermination mechanism^21^. ZTP riboswitch aptamers comprise two subdomains (P1/P2 and P3) that are associated by a pseudoknot and make extensive ZMP-dependent tertiary contacts^21, 26–28^ (Figure 3a, q). ZMP binding is mediated by nucleotides in L3 and by Mg^2+^-mediated recognition of the Z nucleobase^26–28^. It was previously determined that the *pfl* riboswitch folding pathway involves the formation of a non-native intermediate hairpin (IH1) and that ZMP binding coordinates tertiary contacts between the P1/P2 and P3 subdomains that render P3 resistant to terminator hairpin nucleation^25^. However, most of the evidence supporting these conclusions was from 5’-proximal reactivity signatures due to the limited resolution of 3’- proximal nucleotides. To validate TECprobe-ML, we confirmed that the SC1 leader did not meaningfully impact *pfl* ZTP riboswitch function, repeated the analysis of *pfl* riboswitch folding using the SHAPE probe benzoyl cyanide (BzCN)^29, 30^, and extended this analysis to include dimethyl sulfate (DMS) probing^31, 32^ (Figure 3b-d and Supplementary Figure 7a, b).

*pfl* riboswitch folding begins with the rearrangement of IH1 (nts 10-23) to form the P1/P2 subdomain^25^. P1 folding was detected as decreased reactivity in the IH1 loop (BzCN, U16, A17; DMS, nts 15-17) and the downstream P1 stem (BzCN, nts 44-47, U49, A50, A56; DMS, A50, A53, C54) from transcript ∼67 to ∼80, as the downstream P1 stem (nts 44-56) emerges from RNAP (Figure 3a and Supplementary Figure 7e-h, m). P2 folding was detected at transcript ∼60 as decreased P2 reactivity (BzCN, U31, G32; DMS, nts 39-41) when the entire P2 hairpin has emerged from RNAP (Supplementary Figure 7i, j). However, the BzCN reactivity of the upstream P2 stem increases when IH1 refolds into P1 at transcript ∼74, suggesting that the flexibility of P2 increases in the context of the folded P1/P2 subdomain (Supplementary Figure 7i). In previous data, the IH1- to-P1 folding transition was only detectable as decreased IH1 loop reactivity^25^.

The P1/P2 and P3 subdomains are connected by a linker composed of a hairpin and an unstructured region^21^ (Figure 3A). Folding of the linker hairpin was detected as decreased reactivity within the upstream (BzCN, U61, A62; DMS, nts 59-62) and downstream (BzCN, A70; DMS, A72) segments of its stem at transcript 93, when the entire hairpin has emerged from RNAP (Supplementary Figure 7k, l).

The *pfl* riboswitch pseudoknot comprises base pairs between J1/2 (nts 23-27) and L3 (nts 89-93)^21^ (Figure 3a). ZMP-independent pseudoknot folding was detected as decreased reactivity in J1/2 (BzCN, nts 24-26; DMS, A24, C25) from transcript ∼104 to ∼110 when nucleotides 89-93 emerge from RNAP (Figure 3e and Supplementary Figure 7c). Pseudoknot folding is coordinated with decreased BzCN reactivity in P2 (nts U31, G32), in the terminal base pair of P1 (A56), and in P1-proximal unpaired nucleotides within the inter-subdomain linker (A77, C78) (Supplementary Figure 7i, m). These previously undetectable^25^ coordinated reactivity changes indicate that, as expected, pseudoknot folding drives global changes in aptamer structure.

P3 hairpin folding was not observable in previous data due to poor resolution of 3’-proximal nucleotides^25^. Consequently, ZMP binding was inferred from reactivity changes in the P1/P2 subdomain even though ZMP primarily interacts with L3 (Figure 3q). In contrast, the current TECprobe-ML data provide a high-resolution map of P3 folding and direct evidence for ZMP binding. P3 folding was detected as decreased reactivity in the upstream P3 stem (BzCN, nts 84-86; DMS, C85, C86) from transcript 114 to 118, after the downstream P3 stem (nts 96-98) has emerged from RNAP (Figure 3f and Supplementary Figure 7d). In coordination with P3 folding, a ZMP-dependent decrease in BzCN reactivity occurred at U94, which forms two hydrogen bonds with the Z nucleobase, and G95, which stacks with the Z nucleobase^26–28^ (Figure 3g). In agreement with the observation that ZMP binding renders P3 resistant to terminator hairpin nucleation^25^, the BzCN reactivity of the terminator nucleation point (C98) was lower with ZMP (Figure 3h). These previously undetectable^25^ reactivity signatures are direct evidence for ZMP binding and the mechanism of ZMP-mediated transcription antitermination.

P3 folding is coordinated with extensive ZMP-dependent reactivity changes in the P1/P2 subdomain, which were observed previously^25^. First, decreased reactivity at A34 and A38 (BzCN A34, A38; DMS A38) corresponds to reduced L2 flexibility and the formation of a conserved A-minor motif^26–28^, respectively (Figure 3i, j). Second, ZMP-dependent decreased reactivity in upstream (BzCN and DMS, A20, C21) and downstream (BzCN, nts 45-47; DMS, A46) P1 nucleotides and in J2/1 (BzCN, A43) corresponds to the stabilization of non-canonical P1 base pairs, which most likely results from the formation of a ribose zipper between P1 and P3^25–28^ (Figure 3k-n). Consistent with the formation of a Hoogsteen/Hoogsteen base pair between G19 and A47^26–28^, which leaves the Watson-Crick face of A47 unpaired, A47 remained reactive to DMS regardless of whether ZMP was present (Figure 3o). Third, a ZMP-dependent decrease in the BzCN reactivity of A29 that is most likely due to stacking with the J1/2-L3 pseudoknot was previously observed^25^. In the current data, this was not detectable by BzCN probing. However, increased DMS reactivity at A29 suggests that ZMP binding places A29 in an unpaired conformation (Figure 3p).

Formation of the terminator hairpin requires disruption of both P3 and the pseudoknot^21^ (Figure 3a). Without ZMP, terminator hairpin nucleation is detected at transcript 123 as increased reactivity in J1/2 (BzCN, nts 24- 26; DMS, A24, C25) and the upstream P3 stem (BzCN, nts 84-86; DMS, C85, C86), which become unpaired when the J1/2-L3 pseudoknot and P3 are broken (Figure 3e, f and Supplementary Figure 7c, d). At transcript 129, decreased DMS reactivity in the upstream P3 stem (C85, C86) is consistent with complete terminator hairpin folding (Supplementary Figure 7d). With ZMP, the BzCN reactivity of the upstream P3 stem (C85, C86) also increases when the terminator hairpin can nucleate at transcript 123, however there is no corresponding increase in the BzCN or DMS reactivity of J1/2, indicating that the pseudoknot is intact (Figure 3e, f and Supplementary Figure 7c). Notably, the DMS reactivity of the upstream P3 stem (nts C85 and C86) remains low in the presence of ZMP, which suggests that P3 remains intact despite its increased flexibility (Supplementary Figure 7d).

Visualization of terminator readthrough transcripts was difficult using Cotranscriptional SHAPE-Seq because the ZTP riboswitch terminator hairpin is a robust structural barrier to reverse transcription^25^. The reverse transcription conditions used for TECprobe-ML facilitated nearly complete terminator bypass and enabled high-resolution data to be obtained for full-length ZTP riboswitch transcripts (Supplementary Figure 3b). As expected, transcripts from terminally roadblocked TECs adopted alternate structures depending on whether ZMP was included in the reaction. Without ZMP, increased reactivity at J1/2 indicates that the pseudoknot was disrupted (Figure 3e and Supplementary Figure 7c). With ZMP, low reactivity at J1/2 indicates that the pseudoknot remained intact, and the persistence of ZMP-dependent reactivity signatures indicates that ZMP remained bound to the aptamer (Figure 3e, g-n, p and Supplementary Figure 7c). BzCN probing of full-length ZTP riboswitch transcripts using a variation of the TECprobe procedure designed to target single transcript lengths (TECprobe-SL) yielded identical results (Supplementary Figure 8).

In sum, TECprobe-ML detected all *pfl* ZTP riboswitch folding events that were detected previously by Cotranscriptional SHAPE-Seq^25^ and facilitated clear visualization of P3 subdomain folding, ZMP-binding, and terminator readthrough transcripts, which was not previously possible.

### Validation: The *B. cereus crcB* fluoride riboswitch

Watters, Strobel et al. previously used Cotranscriptional SHAPE-Seq to map the folding pathway of the *Bce crcB* fluoride riboswitch^14^, which controls gene expression in response to fluoride using a transcription antitermination mechanism^22^. The fluoride aptamer comprises an H-type pseudoknot and long-range reversed Watson-Crick and Hoogsteen A-U base pairs, and coordinates three Mg^2+^ ions which in turn coordinate the fluoride anion^33^ (Figure 4a, q). The previous analysis of *crcB* riboswitch folding found that fluoride binding stabilizes the pseudoknot, promotes formation of the reversed Watson-Crick A-U pair, reduces the overall flexibility of the aptamer, delays terminator hairpin nucleation, and blocks propagation of terminator base pairs^14^. To further validate TECprobe-ML, we confirmed that the SC1 leader did not meaningfully impact *crcB* fluoride riboswitch function, repeated the analysis of the *crcB* riboswitch using BzCN, and extended this analysis to include DMS probing (Figure 4b-d and Supplementary Figure 9a, b).

**Figure 4.**
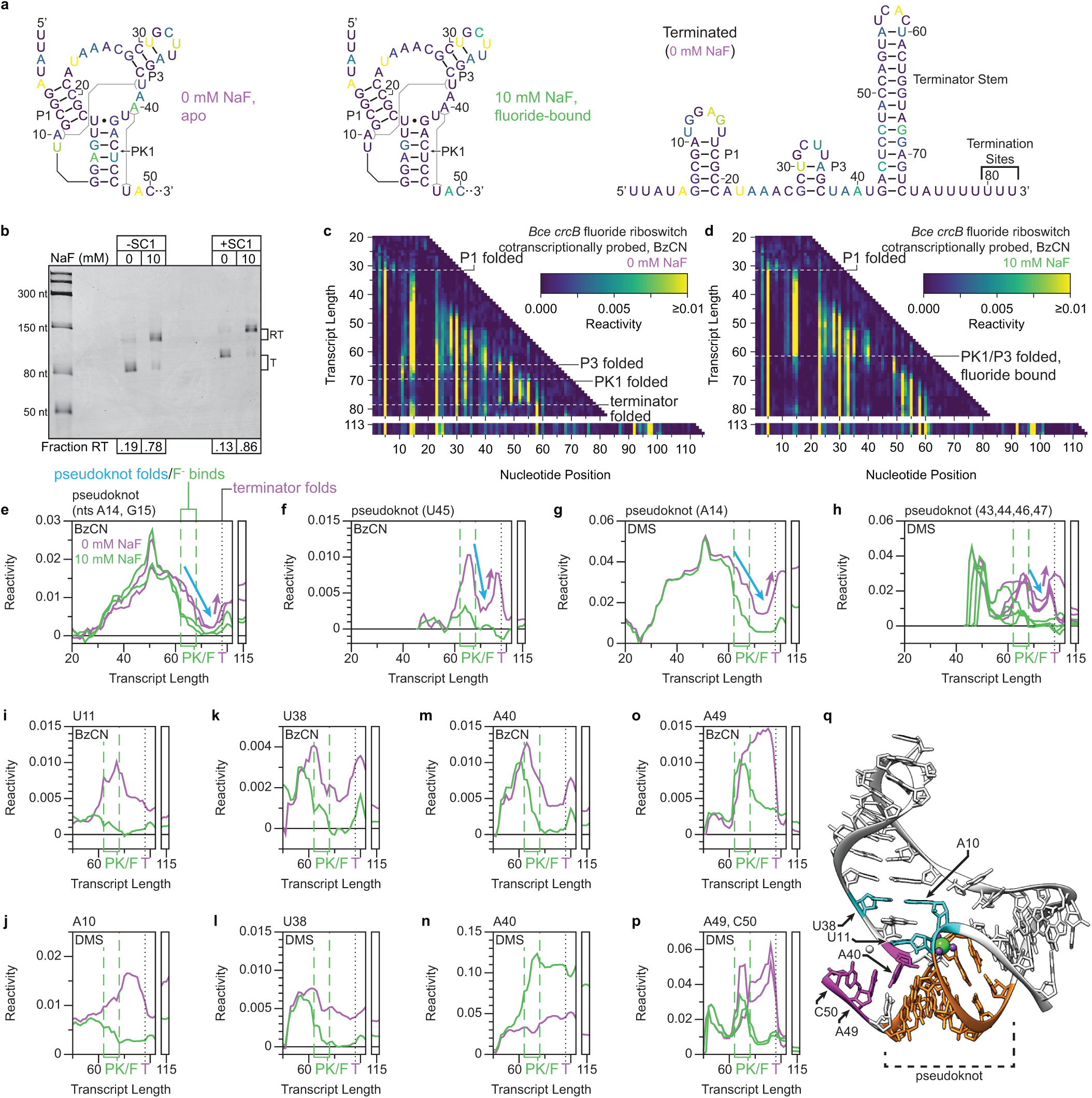
Cotranscriptional folding transitions of the *B. cereus crcB* fluoride riboswitch. **a**, Secondary structures of *crcB* fluoride riboswitch folding intermediates colored by BzCN reactivity (apo/fluoride-bound aptamer: 68 nt; terminated: 120 nt). **b**, Single-round transcription termination assays for the *crcB* fluoride riboswitch with or without SC1. Fraction readthrough is the average of two independent replicates. **c,d**, TECprobe-ML BzCN reactivity matrices for the *crcB* fluoride riboswitch with 0 mM and 10 mM NaF. Transcripts 83-112, which were not enriched, are excluded. Data are from two independent replicates that were concatenated and analyzed together. Reactivity is shown as background-subtracted mutation rate. **e-p**, Transcript length-dependent reactivity changes showing folding transitions. DMS data are from Supplementary Figure 9a, b. Green dashed lines indicate pseudoknot folding. The black dotted line indicates terminator folding. **q**, *Thermotoga petrophila* fluoride riboswitch structure (PDB 4ENC)^33^. Pseudoknot (orange), A10-U38 base-pair-associated (cyan), and A40-U48 linchpin base-pair-associated (magenta) nucleotides are highlighted. F^-^ and Mg^2+^ are green and purple, respectively. BzCN, benzoyl cyanide; PK, pseudoknot; SC1, structure cassette 1; RT, readthrough; T, terminated; DMS, dimethyl sulfate.

The *crcB* riboswitch pseudoknot comprises base pairs between nucleotides 12-17 of L1 and nucleotides 42- 47^22^ (Figure 4a). Pseudoknot folding was detected as decreased reactivity in both L1 (BzCN, A14, G15; DMS, A14) and nucleotides 43-47 (BzCN, U45; DMS, A43, C44, C45, C46) beginning at transcript ∼59, when nucleotides 42-47 are emerging from RNAP (Figure 4e-h). With fluoride, pseudoknot reactivity decreases abruptly from transcript ∼59 to ∼61 and reaches a lower minimum reactivity at transcript ∼70, when the pseudoknot has fully folded, than in the absence of fluoride (Figure 4e-h). This is consistent with the snap-lock model for fluoride aptamer stabilization, in which the apo aptamer explores multiple states dynamically, but the holo aptamer exists in a stably docked state^34^. Previous data did not resolve the fluoride-dependent nature pseudoknot folding and relied solely on reactivity changes within L1 to track pseudoknot folding because of poor resolution in 3’-proximal nucleotides^14^.

Formation of the *crcB* aptamer pseudoknot is coordinated with numerous fluoride-dependent reactivity changes. First, the reactivity of L1 nucleotides that are not involved in pseudoknot base pairs (BzCN, U11; DMS, A10) increases upon pseudoknot folding when fluoride is absent, but decreases when fluoride is present (Figure 4i, j). The lower reactivity of A10 and U11 in the presence of fluoride most likely corresponds to the formation of a reversed Watson-Crick pair between A10 and U38^33^ which extends the P3 stack through U11, and was detected previously^14^ (Figure 4i, j, q). Consistent with this interpretation, the reactivity of U38 also decreases upon pseudoknot folding when fluoride is present (Figure 4k, l). Second, A40 undergoes a fluoride-dependent decrease in BzCN and a fluoride-dependent increase DMS reactivity (Figure 4m, n). This is consistent with the formation of a Hoogsteen A-U pair with U48^33^, known as the linchpin base-pair, which locks the fluoride-bound aptamer into a stably docked state^35^ (Figure 4q). In addition, the reactivity of A49 and C50 (BzCN, A49; DMS, A49, C50), which stack with the linchpin base pair^33^, decrease upon pseudoknot folding in the presence of fluoride (Figure 4o-q). These signatures of linchpin base pair formation were not previously detectable^14^. Third, within J1/3, the reactivity of nucleotides A22 and U23 increases in response to fluoride binding, whereas the reactivity of nucleotides A24 and A25 decreases (Supplementary Figure 9c-g). The fluoride-dependent increase in A22 reactivity was detected in previous data^14^. Fourth, a fluoride-dependent decrease in the DMS reactivity of P3 (C29, C37) indicates that P3 is more stably base-paired in the fluoride-bound aptamer (Supplementary Figure 9h).

It was previously observed that the fluoride-bound *crcB* aptamer delays terminator hairpin nucleation until after RNAP has bypassed the termination sites (+80 to +82) and blocks propagation of terminator base pairs when nucleation eventually occurs^14^. Both fluoride-dependent delayed terminator hairpin formation (Supplementary Figure 9i-o) and inhibition of terminator base pair propagation (Figure 4e-h) were detectable in the current TECprobe-ML data.

In sum, TECprobe-ML detected all *crcB* fluoride riboswitch folding events that were observed previously by Cotranscriptional SHAPE-Seq^14^, and detected several signatures of linchpin base pair formation, which was not previously possible.

### Analysis of ppGpp riboswitch folding using TECprobe-ML

ppGpp riboswitches regulate the expression of genes associated with branched chain amino acid biosynthesis and transport in response to the alarmone ppGpp^23^ and are a member of the *ykkC* aptamer family, which also includes guanidine^36^ and phosphoribosyl pyrophosphate riboswitches^37^ (Figure 5a, b). The ppGpp aptamer comprises two helical stacks (P2-P3 and P1-P4) that are connected by a four-way junction and are associated by long-range contacts between P2 and P4^38, 39^. ppGpp recognition is mediated primarily by the P1-P4 helix and involves direct recognition of the G nucleobase by a conserved C in J3-4^38, 39^. While the overall architecture of ppGpp aptamers and the mechanism of ligand recognition have been established, the mechanism of ppGpp aptamer folding and ppGpp-dependence of aptamer structures remain unresolved. As was the case for the ZTP and fluoride riboswitches above, the SC1 leader hairpin did not meaningfully impact the function of the *Cba* ppGpp riboswitch (Figure 5c). BzCN and DMS probing of ppGpp riboswitch folding intermediates using TECprobe-ML identified several transient structures that occur early in the ppGpp aptamer folding pathway and established which elements of the holo ppGpp aptamer structure are ligand-dependent (Figure 5d-g and Supplementary Figure 10a, b).

**Figure 5.**
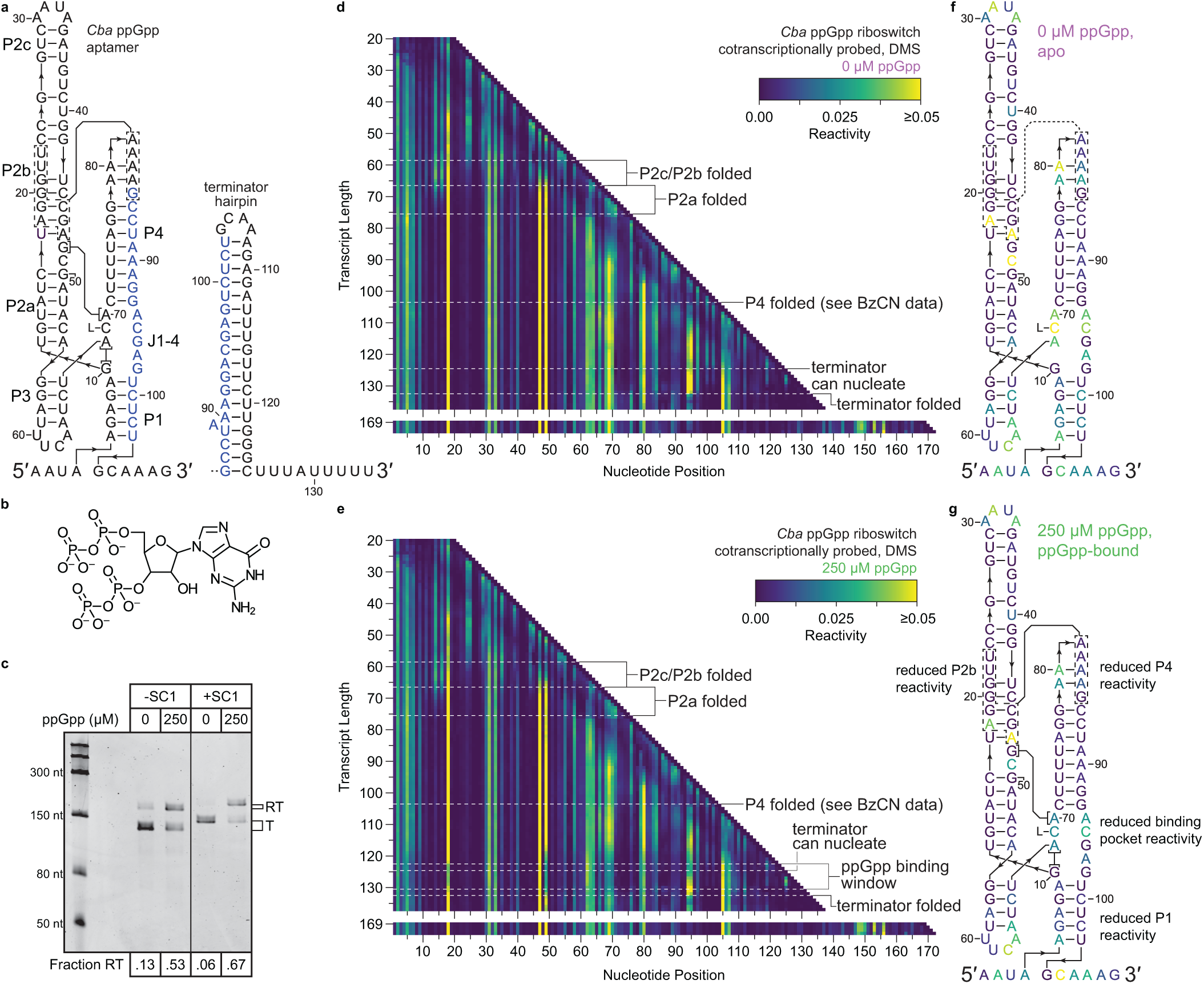
Cotranscriptional chemical probing of the *C. bacterium* ppGpp riboswitch. **a**, Secondary structures of the *Cba* ppGpp aptamer (left) and terminator hairpin (right). Nucleotides present in both the aptamer and terminator hairpin are colored blue, interacting P4 and P2b nucleotides are shown by dashed boxes, and the nucleobase of ppGpp is shown as L. **b**, Chemical structure of ppGpp. **c**, Single-round transcription termination assays for the *Cba* ppGpp riboswitch with or without SC1. Fraction readthrough is the average of four independent replicates. **d**,**e**, TECprobe-ML DMS reactivity matrices for the *Cba* ppGpp riboswitch with 0 μM and 250 μM ppGpp. Transcripts 138-168, which were not enriched, are excluded. Data are from two independent replicates that were concatenated and analyzed together. Reactivity is shown as background-subtracted mutation rate. **f**,**g**, *Cba* ppGpp aptamer secondary structures colored by DMS reactivity from transcript 123 in the absence (**f**) and presence (**g**) of ppGpp. SC1, structure cassette 1; RT, readthrough; T, terminated; DMS, dimethyl sulfate; BzCN, benzoyl cyanide.

### Non-native folding intermediates precede P2 folding

The P2 helix of the *Cba* ppGpp aptamer spans nucleotides 11-55 (Figure 5a). Consequently, P2 cannot fold completely until the nascent transcript is at least 65-70 nt long, and most or all P2 nucleotides have emerged from RNAP. The clearest delineation of early ppGpp aptamer folding is observed when intermediate transcripts are equilibrated prior to BzCN probing, which reduces structural heterogeneity and precisely defines when transient structures become thermodynamically favorable (Supplementary Figure 11a, b). Two intermediate structures were identified: The first intermediate hairpin (IH1) comprises base pairs between nucleotides 7-9 and 17-15; the IH1 stem is non-reactive, whereas nucleotides U11 and A14 of the IH1 loop are reactive to BzCN (Supplementary Figure 11c). The second intermediate hairpin (IH2) comprises base pairs between nucleotides 7-21 and 40-28 with two asymmetrical internal bulges and is mutually exclusive with IH1 (Supplementary Figure 11d). The primary evidence for IH2 is that the reactivity of A14, which is paired in IH2 but not in IH1, decreases when the transcript is 37 nt long and IH2 becomes thermodynamically favorable (Supplementary Figure 11g). Similarly, the reactivity of A14 increases in coordination with decreased reactivity at G19 (IH2 bulge) and U22 (IH2 loop) and increased reactivity at A33 (P2 loop) when the transcript is 45 nt long, which correlates with P2c and P2b folding and disrupting IH2 so that IH1 can fold again (Supplementary Figure 11e, g). The reactivity of A14 and U11 then decreases when the transcript is 53 and 56 nts long, respectively, as each nucleotide becomes paired in P2a (Supplementary Figure 11f, g).

Similar folding transitions were detected in cotranscriptional BzCN and DMS probing data, however, the intermediate structures that occur before P2 folding were less clearly delineated. Formation of P2c was detected as increased BzCN reactivity at A33, which are located in the P2 loop, from transcripts ∼54 to ∼67 in coordination with decreased DMS reactivity at C29, which pairs with G34 in P2c (Figure 6d, f, g, k). P2b folding is coordinated with P2c folding and is detected as decreased BzCN reactivity at G19 and U22, which are not paired in IH1 or IH2, but become constrained in P2b when G19 pairs with C45 and U22 stacks with G21 and G37^38, 39^ (Figure 6a-d, f). The DMS reactivity of C24 and C25, which are unpaired in IH2 and paired in P2b, decreased across the same transcript lengths (Figure 6g-k). In coordination with P2b folding, the reactivity of A14, which is unpaired in IH1 and paired in IH2, increased (Figure 6f, g). Similarly, the BzCN reactivity of U11 in the IH1 loop and the DMS reactivity of C16, which is part of a 3 bp stem in IH1 and a 6 bp stem in IH2, increased to a lesser degree (Figure 6f, g). Because P2b is mutually exclusive with IH2 and reduces the extent to which IH1 can interact with unpaired nucleotides in the nascent transcript, increased IH1 loop reactivity upon P2b folding is likely caused by the collapse of a heterogenous pool of structures into a single folding pathway. These early folding intermediates likely include IH1 and IH2, but may also include other non-native structures that were not clearly identified in equilibrium chemical probing experiments (Figure 6b, c, i, j). P2a folding is then observed as decreased reactivity at U11, A14, and C16 from transcript ∼67 to ∼76 as these nucleotides become paired in P2a, and increased DMS reactivity at A18, which forms a Hoogsteen/sugar edge pair with G46 in P2b^38, 39^ (Figure 6 e-g, l).

**Figure 6.**
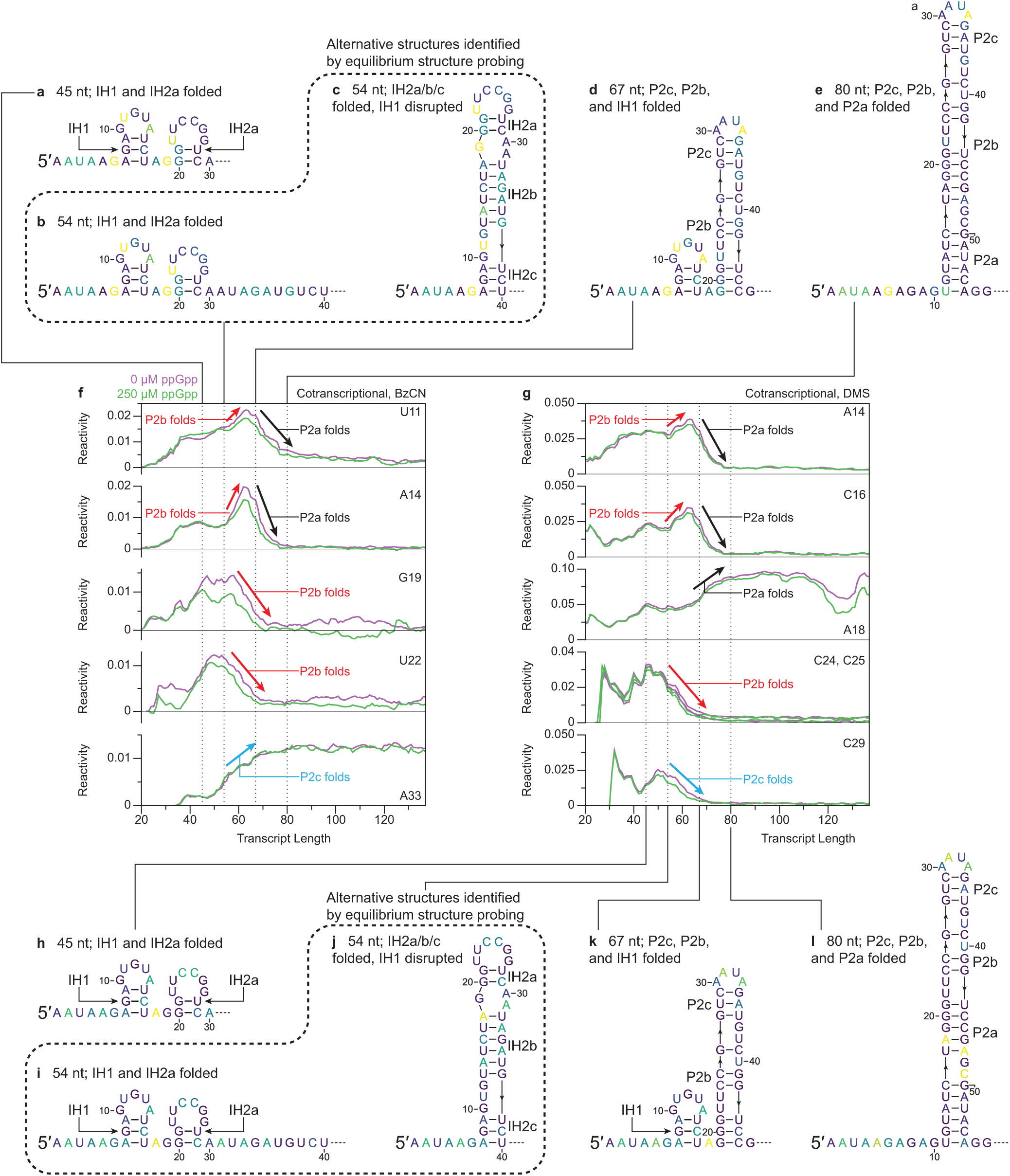
Early folding intermediates of the *C*. *bacterium* ppGpp riboswitch. **a-e**, Secondary structures of *Cba* ppGpp riboswitch folding intermediates containing IH1 and IH2a (**a**, **b**), IH2a/b/c (**c**), IH1 and P2b/c (**d**), and P2a/b/c (**e**) colored by the BzCN reactivity of the indicated transcript in the absence of ppGpp. **f**,**g**, Plots showing transcript length-dependent BzCN (**f**) or DMS (**g**) reactivity changes that occur during P2 hairpin folding. BzCN reactivity is from Supplementary Figure 10a, b. DMS reactivity is from Figure 5d, e. Arrows indicate reactivity changes associated with P2a (black), P2b (red), and P2c (blue) folding. **h-l**, As in **a**-**e**, except that secondary structures are colored by the DMS reactivity of the indicated transcript lengths. Reactivity is shown as raw background-subtracted mutation rate. BzCN, benzoyl cyanide; DMS, dimethyl sulfate.

### ppGpp binding stabilizes a pre-formed ligand binding pocket

ppGpp binding was detected at transcript ∼123 as decreased DMS reactivity in nucleotides proximal to the ppGpp-binding pocket (Figure 7a, b). The most direct signature of ppGpp binding is decreased DMS reactivity at C69, which forms a Watson-Crick pair with the G nucleobase of ppGpp^38, 39^ (Figure 7b, c). The ppGpp-dependent formation the G10-A68 pair, which stacks with the C69-ppGpp pair^38, 39^, was detected as decreased DMS reactivity at A68 (Figure 7b, d). Similarly, the ppGpp-dependent formation of contacts between G48 and A70, which stacks on the opposite face of the C69-ppGpp pair^38, 39^, was detected as decreased DMS reactivity at A70 (Figure 7b, d). The DMS reactivity of C49, which stacks with G48 and coordinates a Mg^2+^ that contacts the 3’ terminal phosphate of ppGpp^38, 39^, and the BzCN reactivity G98, which directly contacts the 5’ terminal phosphate of ppGpp^38^, also decrease upon ppGpp binding (Figure 7b, e and Supplementary Figure 10f). In contrast to the ppGpp-dependent changes in the DMS reactivity of nucleotides 68-70, the BzCN reactivity of nucleotides 68-70 decreases at transcript ∼115 independent of ppGpp binding (Supplementary Figure 10c-e). Given that BzCN measures backbone flexibility and DMS measures Watson-Crick face accessibility, this suggests that folding of the apo aptamer partially organizes the ligand binding pocket, but that ppGpp binding is required to establish the A68-G10 and A70-G48 pairs. The DMS reactivity of nucleotides A94 and C95 within J4-1, which are proximal to the ppGpp binding pocket^38, 39^, also decreases upon ppGpp binding (Figure 7b, f, g). All ppGpp-dependent reactivity changes described above and in the sections below were abolished by the C69A point mutation, which disrupts ppGpp binding^23^ (Figure 7c-g and Supplementary Figure 12).

**Figure 7.**
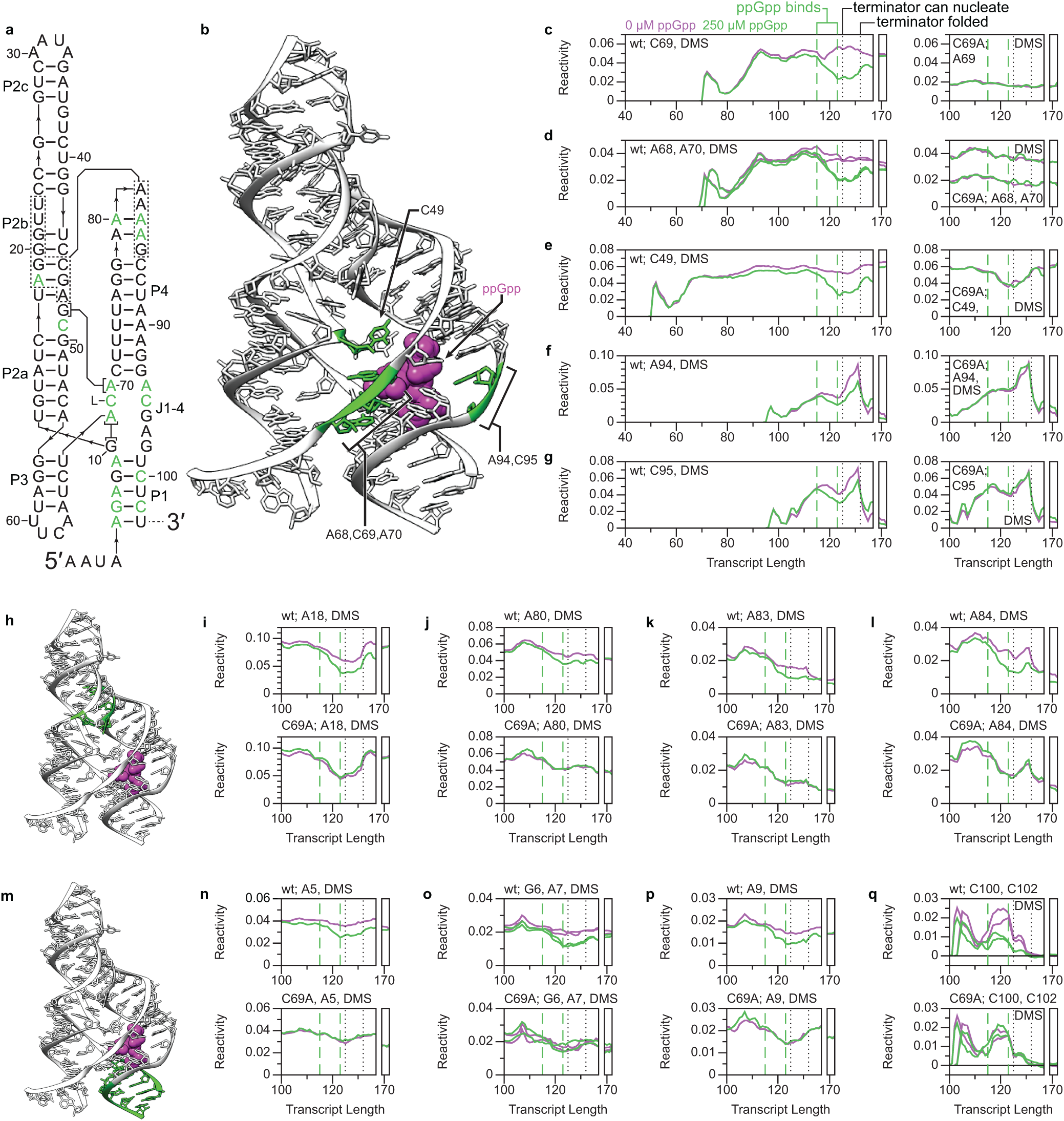
Ligand-dependent changes in ppGpp aptamer structure. **a**, *Cba* ppGpp aptamer secondary structure. Nucleotides colored green exhibit reduced reactivity upon ppGpp binding. **b,h,m,** *Thermoanaerobacter mathranii* PRPP riboswitch G96A (ppGpp-binding) variant structure (PDB 6CK4)^38^ highlighted to indicate ppGpp-responsive nucleotides in the ligand binding pocket (**b**) and within P2b-P4 contacts (**h**), and ppGpp-responsive base pairs in P1 (**m**). **c-g**, **i-l**, **n-q**, Plots of transcript length-dependent reactivity changes for ppGpp-responsive nucleotides in the ligand binding pocket (**c-g**), within P2b-P4 contacts (**i-l**), and within P1 (**n-q**) for the WT *Cba* ppGpp riboswitch and C69A variant. Green dashed lines indicate the bounds of the apo-to-ppGpp-bound folding transition. Black dotted lines indicate positions where the terminator begins to nucleate and is folded. Data are from Figure 5d, e and Supplementary Figure 12a, b. Reactivity is shown as background-subtracted mutation rate. DMS, dimethyl sulfate.

### ppGpp binding stabilizes a distal tertiary interaction

Crystal structures of ppGpp aptamers revealed that a stretch of purines in P4 forms long-range interactions with the minor groove of P2b^38, 39^ (Figure 7a, h). Independent of ppGpp binding, the DMS reactivity of A18, which is the only DMS-reactive (A or C) nucleotide in P2b with an unpaired Watson-Crick face^38, 39^, and P4 (nts A80, A83, A84) decreases at transcript ∼108 (Figure 7i-l). This is consistent with the formation of long-range P2b-P4 contacts when P4 has fully emerged from RNAP. In coordination with the signatures of ppGpp binding described above, these P2b and P4 nucleotides undergo a further ppGpp-dependent decrease in DMS reactivity at transcript ∼115 after the ppGpp aptamer has fully emerged from RNAP (Figure 7i-l). It is important to note that the reactivity changes observed in P4 cannot confidently be attributed to specific nucleotides because P4 contains 6 consecutive A nucleotides, which prevents precise cDNA mutation mapping^20^.

Nonetheless, the observation of sequential ppGpp-independent and ppGpp-dependent decreases in the DMS reactivity of P2b and P4 suggests that ppGpp binding stabilizes the interaction between the two helices despite the distance of the P2b-P4 contacts from the ligand binding pocket. Consistent with this interpretation, the ppGpp binding-deficient C69A mutant undergoes the ppGpp-independent decrease in DMS reactivity at transcript ∼108 but not the ppGpp-dependent decrease in DMS reactivity at transcript ∼115 (Figure 7i-l).

### ppGpp binding stabilizes P1 base pairs against terminator hairpin folding

It has been proposed that ppGpp binding promotes transcription antitermination by stabilizing P1, which blocks propagation of terminator base pairs^38, 39^. In agreement with this model, we observed a ppGpp-dependent decrease in the reactivity of every P1 base pair (Figure 7a, m-q and Supplementary Figure 10g, h). The folded ppGpp riboswitch terminator hairpin was detected in cotranscriptional and equilibrium BzCN probing experiments as a low reactivity region spanning nucleotides 85 to 125 (Supplementary Figure 13a-d). Within this region, A107, which corresponds to the ‘A’ within the apical GNRA tetraloop, and A89, which is predicted to be a single nucleotide bulge, were reactive (Supplementary Figure 13a-f). In the absence of ppGpp, terminator nucleation was detected as decreased DMS reactivity at C100 and C102, which form base pairs near the apical loop of the terminator, at transcript ∼125 when ∼4 terminator base pairs can form outside of RNAP (Figure 7q). In the presence of ppGpp, terminator nucleation at transcript ∼125 was coordinated with weakly increased BzCN reactivity in P2b and P4 even though all observed ppGpp-dependent differences in DMS reactivity were maintained (Figure 7 and Supplementary Figures 10b and 13g). It is unclear whether this change in nucleotide flexibility is due to a sub-population of ppGpp aptamers being disrupted by terminator nucleation, or if emergence of the terminator loop and downstream stem nucleotides from RNAP perturbs the ppGpp aptamer without disrupting it. When RNAP reaches the primary termination sites at positions +132/133, completion of the terminator stem is observed as increased BzCN reactivity at the A89 bulge and a partial reduction of ppGpp-dependent differences in DMS reactivity as some ppGpp-bound aptamers are disrupted by the terminator hairpin (Figure 7 and Supplementary Figure 13f). In the ZTP and fluoride riboswitches described above, ligand-dependent structures persisted after RNAP had bypassed the transcription termination site. In contrast, ppGpp-dependent reactivity signatures were not detectable in transcripts that accumulated at the terminal transcription roadblock downstream of the termination site (Figure 7 and Supplementary Figure 13h-l). One possible explanation for this effect is that dissociation of ppGpp from the aptamer allows the terminator to nucleate.

## Discussion

We have demonstrated that TECprobe-ML detects biologically meaningful RNA folding intermediates at nucleotide resolution and delineates when coordinated structural rearrangements can occur within a cotranscriptional RNA folding pathway. TECprobe-ML simplifies and standardizes cotranscriptional RNA structure probing experiments by using MaP^20^ to encode RNA structural information in cDNA, and by structuring cDNA as an amplicon using a 5’ structure cassette. This approach eliminates the need for size selection procedures that can cause information loss near transcript 3’ ends and ensures even sequencing coverage across the length of each transcript. The benefits of this approach are evident in our reinvestigation of ZTP and fluoride riboswitch folding and our analysis of ppGpp riboswitch folding: in all cases, improved resolution of 3’ proximal RNA structures enabled the detection of cotranscriptional folding events in which coordinated reactivity changes spanned the entire transcript. In combination with complementary biophysical approaches^8–13^, the ability to resolve coordinated structural changes by cotranscriptional RNA chemical probing will likely aid efforts to predict RNA folding pathways^40–46^.

There are two potential drawbacks to structuring cDNA as an amplicon in the TECprobe-ML procedure: first, structuring cDNA as an amplicon requires the inclusion of a 5’ structure cassette in the target RNA sequence, which could potentially influence RNA folding in some cases. However, the use of structure cassettes in RNA chemical probing experiments is well-validated^47^ and did not interfere with the function or folding of the three riboswitches that were investigated in this study. Second, obtaining complete coverage of cDNA requires the use of sequencing reads that span the length of the longest intermediate transcript in the library. Consequently, the upper bound for target RNA length is currently ∼440 nt, which accounts for a constant primer binding site downstream of the target RNA sequence, the 5’ structure cassette, barcoding modified and untreated samples, and a buffer to accommodate insertions in the cDNA that can occur during error-prone reverse transcription. At the time of writing, the maximum target RNA length we have tested is 273 nt.

Like all cotranscriptional RNA structure probing methods in which nascent RNA is chemically probed in the context of a roadblocked RNAP, TECprobe-ML has two primary limitations. First, TECprobe-ML does not measure true cotranscriptional folding because transcription must be artificially halted at a roadblock before chemical probing can be performed. Given that base pair formation occurs on a microsecond timescale^4^, halting transcription could permit local equilibration of the cotranscriptionally folded RNA. Second, although it is useful to visualize reactivity data from cotranscriptional RNA structure probing experiments as a continuous trajectory as we have done above, the profile of each intermediate transcript is technically an end-point measurement. Therefore, current experiments cannot definitively link the intermediate structures that are observed within a cotranscriptional folding pathway. Notably, these limitations are complementary to those of biophysical approaches, which have high temporal resolution and track RNA folding continuously, but which do not resolve RNA secondary or tertiary structure directly.

## Materials and Methods

### Oligonucleotides

All oligonucleotides were purchased from Integrated DNA Technologies. A detailed description of all oligonucleotides including sequence, modifications, and purifications is presented in Supplementary Table 1.

### Proteins

Q5 High-Fidelity DNA Polymerase, Vent (exo-) DNA polymerase, *Sulfolobus* DNA Polymerase IV, *E*. *coli* RNA Polymerase holoenzyme, Mth RNA Ligase (as part of the 5’ DNA Adenylation kit), T4 RNA Ligase 2 truncated KQ, ET SSB, RNase H, and RNase If were purchased from New England Biolabs. TURBO DNase, SuperaseIN, SuperScript II, SuperScript III, and BSA were purchased from ThermoFisher. Streptavidin was purchased from Promega. NusA was a gift from J. Roberts (Cornell University)

### DNA template purification

Supplementary Table 2 provides details for the oligonucleotides and processing steps used for every DNA template preparation in this work. Supplementary Table 3 provides DNA template sequences.

5’ biotinylated DNA that was used for transcription antitermination assays and as a PCR template when preparing modified DNA templates below was PCR amplified from plasmid DNA using Q5 High-Fidelity DNA Polymerase (New England Biolabs) and primers HP4_5bio.R and PRA1_NoMod.F (for ZTP and fluoride templates) or PRA1_shrt.F (for ppGpp templates) (Supplementary Table 1) in a 300 μl PCR, and purified by UV-free agarose gel extraction as described previously^48^; PCR was performed as 100 μl reactions. A step-by-step protocol for this procedure is available^49^.

Randomly biotinylated DNA templates for TECprobe-ML experiments were PCR amplified from a 5’ biotinylated linear DNA template using Vent (exo-) DNA polymerase (New England Biolabs) and primers HP4_5bio.R and PRA1_NoMod.F (for ZTP and fluoride templates) or PRA1_shrt.F (for ppGpp templates) (Supplementary Table 1). 200 μl PCRs contained 1X ThermoPol Buffer (New England Biolabs), 250 nM PRA1_NoMod.F or PRA1_shrt.F, 250 nM HP4_5bio.R, 20 pM template DNA, 0.02 U/μl Vent (exo-) DNA polymerase, 200 μM dNTP Solution Mix (New England Biolabs), and a concentration of biotin-11-dNTPs (PerkinElmer, Biotium) that favored the incorporation of ∼2 biotin modifications in the transcribed region of each DNA template (which varied based on the DNA template sequence and was calculated as described previously^15^). PCR was performed as two 100 μl reactions in thin-walled tubes using the following thermal cycler protocol with a heated lid set to 105°C: 95°C for 3 min, [95°C for 20 s, 58°C for 30 s, 72°C for 30 s] x 30 cycles, 72°C for 5 min, hold at 12°C. PCR products were purified as described below in the section *SPRI bead purification of DNA*, eluted into 50 μl of 10 mM Tris-HCl (pH 8.0), and quantified using the Qubit dsDNA Broad Range Assay Kit (Invitrogen) with a Qubit 4 Fluorometer (Invitrogen). A step-by-step protocol for this procedure is available^49^.

DNA templates for TECprobe-SL experiments, which contained an internal etheno-dA stall site, were prepared using a variation of a previously described^48, 50^ procedure for preparing DNA that contains a chemically-encoded transcription stall site. In this variation of the procedure, PCR products were purified using SPRI beads instead of by column purification. Briefly, DNA with an internal etheno-dA stall site was PCR amplified from an unmodified linear DNA template using Q5 High-Fidelity DNA Polymerase and primers PRA1_NoMod.F and dRP1iEthDA.R in a 300μl PCR, as described previously^48, 50^; PCR was performed as 100 μl reactions.

PCR products were purified as described below in the section *SPRI bead purification of DNA* and eluted into 100 μl of 10 mM Tris-HCl (pH 8.0) per 300 μl PCR. Translesion DNA synthesis using *Sulfolobus* DNA polymerase IV (New England Biolabs) was performed as two 100 μl reactions as described previously^48, 50^.

After translesion DNA synthesis, DNA templates were purified as described below in the section *SPRI bead purification of DNA*, eluted into 75 μl of 10 mM Tris-HCl (pH 8.0), and quantified using the Qubit dsDNA Broad Range Assay Kit with a Qubit 4 Fluorometer. DNA templates for experiments that evaluated reverse transcription additives were prepared as above except dRP1iBio.R, which contains an internal biotin-TEG modification, was used.

### SPRI bead purification of DNA

SPRI beads were prepared in-house using the ‘DNA Buffer’ variation of the procedure by Jolivet and Foley^51^. Samples were mixed with an equal volume of SPRI beads, incubated at room temperature for 5 min, and placed on a magnetic stand for 3 min so that the beads collected on the tube wall. The supernatant was aspirated and discarded, and the beads were washed twice by adding a volume of 70% ethanol at least 200 μl greater than the combined volume of the sample and SPRI beads to the tube without disturbing the bead pellet while it remained on the magnetic stand. The samples were incubated at room temperature for 1 min before aspirating and discarding the supernatant. Residual ethanol was evaporated by placing the open microcentrifuge tube in a 37°C dry bath for ∼15 s with care taken to ensure that the beads did not dry out.

Purified DNA templates were eluted by resuspending the beads in a variable amount of 10 mM Tris-HCl (pH 8.0) (depending on the procedure, details are in each relevant section), allowing the samples to sit undisturbed for 3 min, placing the sample on a magnetic stand for 1 min so that the beads collected on the tube wall, and transferring the supernatant, which contained purified DNA, into a screw-cap tube with an O-ring.

### Transcription antitermination assays

Single-round *in vitro* transcription using terminally biotinylated template DNA was performed as described for TECprobe-ML below in the section *Single-round in vitro transcription for TECprobe-ML and TECprobe-SL* except that the reaction volume was 25 μl for ZTP and ppGpp templates or 50 μl for fluoride templates, 100 nM streptavidin (Promega) was included in the master mix, and the transcription reactions were stopped by adding 75 μl (for ZTP and ppGpp templates) or 150 μl (for fluoride templates) of TRIzol LS without performing RNA chemical probing. RNA was purified as described in the section *RNA purification*, except that volumes of chloroform and isopropanol were scaled to account for the doubled reaction volume for fluoride templates and all samples were dissolved in 6 (ppGpp samples) or 15 μl (ZTP and fluoride samples) of formamide loading dye (90% (v/v) deionized formamide, 1X transcription buffer (defined below), 0.05% (w/v) bromophenol blue, 0.05% (w/v) xylene cyanol FF), and subsequently analyzed as described in the section *Denaturing Urea-PAGE*.

### Denaturing Urea-PAGE

Samples in formamide loading dye were heated at 95°C for 5 min, and snap-cooled on ice for 2 min. Denaturing Urea-PAGE was performed using 8 or 10% gels prepared using the SequaGel UreaGel 19:1 Denaturing Gel System (National Diagnostics) for a Mini-PROTEAN Tetra Vertical Electrophoresis Cell as described previously^52^. Gels were stained with SYBR Gold Nucleic Acid Stain (Invitrogen) and scanned using an Azure Biosystems Sapphire Biomolecular Imager as described previously^52^.

### TECprobe-ML and TECprobe-SL

We previously described the theory and best practices for cotranscriptional RNA structure probing experiments, and detailed procedures for randomly biotinylated DNA template preparation, cotranscriptional RNA chemical probing, RNA purification, and 3’ adapter ligation^49^.

#### Single-round in vitro transcription for TECprobe-ML and TECprobe-SL

All single-round *in vitro* transcription reactions for TECprobe-ML experiments were performed as 60 μl reactions containing 1X Transcription Buffer [20 mM Tris-HCl (pH 8.0), 50 mM KCl, 1 mM dithiothreitol (DTT), and 0.1 mM EDTA], 0.1 mg/ml Molecular Biology-Grade BSA (Invitrogen), 100 μM high-purity NTPS (Cytiva), 10 nM randomly biotinylated template DNA, and 0.024 U/μl *E. coli* RNA polymerase holoenzyme (New England Biolabs). Single-round *in vitro* transcription reactions for TECprobe-SL experiments were performed as 60 μl reactions containing the same reagents as TECprobe-ML experiments except that the reactions used a 10 nM template DNA that contains an internal etheno-dA stall site downstream of a *Cbe pfl* ZTP riboswitch variant in which the poly-U tract was removed, and either 100 or 500 μM high-purity NTPs, as indicated.

Transcription reactions for experiments where DMS was used to probe RNA structures contained additional Tris-HCl (pH 8.0) at final concentration of 100 mM to minimize pH changes during the chemical probing reaction. At the time of preparation, each TECprobe-ML reaction was 48 μl due to the omission of 10X (1 μM) streptavidin (Promega) and 10X Start Solution [100 mM MgCl2, 100 μg/ml rifampicin (Gold Biotechnology)] from the reaction. Each TECprobe-SL reaction was 54 μl due to the omission of 10X Start Solution from the reaction.

The composition of the single-round *in vitro* transcription master mix varied depending on the riboswitch system that was assessed. *In vitro* transcription reactions for the *Cbe pfl* ZTP riboswitch contained 2% (v/v) DMSO and, when present, 1 mM ZMP (Sigma-Aldrich). *In vitro* transcription reactions for the *Bce crcB* fluoride riboswitch that were performed in the presence of fluoride contained 10 mM NaF (Sigma-Aldrich). *In vitro* transcription reactions for the *Cba* ppGpp riboswitch and C69A ppGpp riboswitch variant contained 500 nM NusA and, when present, 250 μM ppGpp (Guanosine-3’,5’-bisdiphosphate) (Jena Bioscience).

Single-round *in vitro* transcription reactions were incubated at 37°C for 10 min to form open promoter complexes. For TECprobe-ML reactions, 6 μl of 1 μM streptavidin was then added for a final concentration of 100 nM streptavidin, and reactions were incubated for an additional 10 min at 37°C; TECprobe-SL reactions did not include streptavidin but were still incubated for a total of 20 min at 37°C. Transcription was initiated by adding 6 μl of 10X Start Solution to the reaction for a final concentration of 10 mM MgCl2 and 10 μg/ml rifampicin. The transcription reaction was incubated at 37°C for 2 min before chemical probing was performed as described below in the section *RNA chemical probing*.

#### RNA chemical probing

Benzoyl cyanide (BzCN) probing was performed by splitting the sample into 25 μl aliquots and mixing with 2.78 μl of 400 mM BzCN (Sigma-Aldrich) dissolved in anhydrous DMSO (Sigma-Aldrich) [(+) sample)] or with anhydrous DMSO [(-) sample]^29, 30^. Given that the BzCN probing reaction is complete within ∼1 s, 75 μl of TRIzol LS reagent (Invitrogen) was added to each sample without any additional incubation time to stop the *in vitro* transcription reaction and the samples were vortexed.

Dimethyl sulfate (DMS) probing was performed by splitting the sample into 25 μl aliquots and mixing with 2.78 μl of 6.5% (v/v) DMS (Sigma-Aldrich) in anhydrous ethanol (Sigma-Aldrich) [(+) sample)] or with anhydrous ethanol [(-) sample] and incubating the samples at 37°C for 5 min. The DMS probing reaction was quenched by adding beta-mercaptoethanol to 2.8 M and incubating the sample at 37°C for 1 min. 75 μl of TRIzol LS reagent was added to each sample to stop the *in vitro* transcription reaction and the samples were vortexed.

#### RNA purification

Samples, which contained 27.78 μl (BzCN probing) or 34.45μl (DMS probing) of the cotranscriptional RNA chemical probing reaction in 75 μl of TRIzol LS, were extracted as follows: 20 μl of chloroform was added to each sample, and the samples were mixed by vortexing and inverting the tube and centrifuged at 18,500 x g and 4°C for 5 min. The aqueous phase was transferred to a new tube and precipitated by adding 1.5 μl of GlycoBlue Coprecipitant (Invitrogen) and 50 μl of ice-cold isopropanol and incubating at room temperature for 15 min. The samples were centrifuged at 18,500 x g and 4°C for 15 min, the supernatant was aspirated and discarded, 500 μl of ice cold 70% ethanol was added to each sample, and the tubes were gently inverted to wash the samples. The samples were centrifuged at 18,500 x g and 4°C for 2 min and the supernatant was aspirated and discarded. The samples were centrifuged again briefly to pull down residual liquid, which was aspirated and discarded. The pellet was then resuspended in 25 μl of 1X TURBO DNase buffer (Invitrogen), mixed with 0.75 μl of TURBO DNase (Invitrogen), and incubated at 37°C for 15 min. 75 μl of TRIzol LS reagent was added to stop the reactions and a second TRIzol extraction was performed as described above, except that the pellet was resuspended in 5 μl of 10% (v/v) DMSO for TECprobe-ML reactions or 25 μl 1X Buffer TM (1X Transcription Buffer, 10 mM MgCl2) for TECprobe-SL reactions. TECprobe-ML sample processing continued with the *RNA 3’ adapter ligation* section immediately below, while TECprobe-SL samples proceeded directly to the *cDNA synthesis and cleanup* section because the RNA 3’ adapter sequence was included in the transcript.

#### RNA 3’ adapter ligation

9N_VRA3 adapter oligonucleotide (Supplementary Table 1) was pre-adenylated with the 5’ DNA Adenylation Kit (New England Biolabs) according to the manufacturer’s protocol at a 5X scale. Briefly, 100 μl of a master mix that contained 1X DNA Adenylation Buffer (New England Biolabs), 100 μM ATP, 5 μM 9N_VRA3 oligo, and 5 μM Mth RNA Ligase (New England Biolabs) was split into two 50 μl aliquots in thin-walled PCR tubes and incubated at 65°C in a thermal cycler with a heated lid set to 105°C for 1 hour. Following the reaction, 150 μl of TRIzol LS reagent was added to each 50 μl reaction and the samples were extracted as described above in the section *RNA purification*, except that reaction volumes were scaled to account for the 50 μl reaction volume (40 μl of chloroform was added to the sample-TRIzol mixture and 100 μl of isopropanol was added during the precipitation step). Samples were pooled by resuspending the pellets from each TRIzol extraction in a single 25 μl volume of TE Buffer (10 mM Tris-HCl (pH 7.5), 0.1 mM EDTA). The concentration of the adenylated oligonucleotide was determined using the Qubit ssDNA Assay Kit (Invitrogen) with a Qubit 4 Fluorometer. The molarity of the linker was calculated using 11,142 g/mol as the molecular weight. The adenylation reaction was assumed to be 100% efficient. The linker was diluted to 0.9 μM and aliquoted for future use; aliquots were used within 3 freeze-thaw cycles.

20 μl RNA 3’ adapter ligation reactions were performed by combining purified RNA in 5 μl of 10% DMSO (v/v) from the *RNA purification* section with 15 μl of an RNA ligation master mix such that the final 20 μl reaction contained purified RNA, 2.5% (v/v) DMSO, 1X T4 RNA Ligase Buffer (New England Biolabs), 0.5 U/μl SuperaseIN (Invitrogen), 15% (w/v) PEG 8000, 45 nM 5’-adenylated 9N_VRA3 adapter, and 5 U/μl T4 RNA Ligase 2, truncated, KQ (New England Biolabs). The samples were mixed by pipetting and incubated at 25°C for 2 hours.

#### SPRI bead purification of RNA

Excess 9N_VRA3 3’ adapter oligonucleotide was depleted using a modified SPRI bead purification that contains isopropanol^53^. 17.5 μl of nuclease-free water and 40 μl of freshly aliquoted anhydrous isopropanol (Sigma-Aldrich) were added to each 20 μl RNA ligation reaction, and the samples were mixed by vortexing. Each sample was then mixed with 22.5 μl of SPRI beads so that the concentration of PEG 8000 was 7.5% (w/v) and the concentration of isopropanol was 40% (v/v) in a sample volume of 100 μl. The samples were incubated at room temperature for 5 min, and placed on a magnetic stand for at least 3 min so that the beads collected on the tube wall. The supernatant was aspirated and discarded, and the beads were washed twice by adding 200 μl of 80% (v/v) ethanol to the tubes without disturbing the bead pellet while it remained on the magnetic stand, incubating the samples at room temperature for 1 min, and aspirating and discarding the supernatant. After discarding the second 80% (v/v) ethanol wash, the sample was briefly spun in a mini centrifuge, and placed back onto a magnetic stand for 1 min to collect the beads on the tube wall. The supernatant was aspirated and discarded and the beads were briefly (<15 s) dried in a 37°C dry bath with the cap of the microcentrifuge tube left open. Purified RNA was eluted by resuspending the beads in 20 μl of 10 mM Tris-HCl (pH 8.0), incubating the sample for 3 min at room temperature, placing the sample on a magnetic stand for 1 min so that the beads collected on the tube wall, and transferring the supernatant into a clean microcentrifuge tube. The eluted RNA was mixed with 11.5 μl of RNase-free water and 40 μl of anhydrous isopropanol. 6 μl of 50% (w/v) PEG 8000 and 22.5 μl of SPRI beads were added to each sample so that the concentration of PEG 8000 was 7.5% (w/v) and the concentration of isopropanol was 40% (v/v) in a sample volume of 100 μl. The RNA was purified as described above a second time, except that the RNA was eluted into 25 μl of 1X Buffer TM.

### cDNA synthesis and cleanup

5 μl of 10 mg/ml Dynabeads MyOne Streptavidin C1 beads (Invitrogen) per sample volume were equilibrated in Buffer TX (1X Transcription Buffer, 0.1% (v/v) Triton X-100) as described previously^52^. The Streptavidin C1 beads were resuspended at a concentration of ∼2 μg/μl in Buffer TX (25 μl per sample volume), distributed into 25 μl aliquots, and stored on ice until use.

The reverse transcription primer, dRP1_5Bio.R (Supplementary Table 1), contains a 5’ biotin modification and anneals to the 9N_VRA3 adapter (TECprobe-ML) or the transcribed VRA3 sequence downstream of the riboswitch (TECprobe-SL). 1 μl of 500 nM dRP1_5Bio.R was added to the purified RNA, and the samples were incubated on a thermal cycler with a heated lid set to 105°C using the ‘RT anneal’ protocol: pre-heat to 70°C, 70°C for 5 min, ramp to 50°C at 0.1°C/s, 50°C for 5 min, ramp to 40°C at 0.1°C/s, 40°C for 5 min, cool to 25°C. Equilibrated Streptavidin C1 beads were placed on a magnetic stand to collect the beads on the tube wall, and the supernatant was removed. The bead pellets were resuspended using the samples (which contained dRP1_5Bio.R oligo annealed to RNA), and incubated at room temperature for 15 min on an end-over-end rotator set to ∼8 rpm. The samples were placed on a magnetic stand to collect the beads on the tube wall, the supernatant was aspirated and discarded, and the beads were resuspended in 19.5 μl of reverse transcription master mix, which omits SuperScript II reverse transcriptase at this time. After the 20 μl reverse transcription reaction was completed by adding 0.5 μl of SuperScript II as described below, the concentration of each reagent that is present in the master mix was: 50 mM Tris-HCl (pH 8.0), 75 mM KCl, 0.5 mM dNTP Solution Mix, 10 mM DTT, 2% (v/v) deionized formamide (Millipore), 10 ng/μl ET SSB (New England Biolabs), 3 mM MnCl2 (Fisher Scientific), 0.1% (v/v) Triton X-100, and 5 U/μl SuperScript II (Invitrogen). As described in the original SHAPE-MaP procedure, it is **crucial** to add MnCl2 stock to the master mix immediately before performing reverse transcription because the manganese will begin to oxidize and precipitate in this solution^54^. Samples were placed on a pre-heated 42°C thermal cycler for 2 min before 0.5 μl of SuperScript II was added to complete the master mix. The reverse transcription reaction was incubated at 42°C for 50 min, and then at 70°C for 15 min to heat inactivate SuperScript II. Samples were cooled to 12°C, 1.25 U of RNase H (New England Biolabs) and 12.5 U of RNase If (New England Biolabs) were added to each sample, and the samples were incubated at 37°C for 20 min and then at 70°C for 20 min to heat inactivate the RNases. The samples were briefly spun down in a mini-centrifuge and placed on a magnetic stand. The supernatant was aspirated and discarded and the bead pellet was washed with 75 μl Storage Buffer (10mM Tris-HCl (pH 8.0) and 0.05% (v/v) Triton X-100). The beads were resuspended in 25 μl of Storage Buffer, transferred to screw-cap tubes with an O-ring, and stored at -20°C.

### Chemical probing of equilibrium refolded intermediate transcripts

Single-round *in vitro* transcription using randomly biotinylated ppGpp template DNA was performed as described for TECprobe-ML in the section *Single-round in vitro transcription for TECprobe-ML and TECprobe- SL* and transcription was stopped by adding 150 μl of TRIzol LS. Intermediate transcripts were then purified as described in the section *RNA purification* except that volumes of chloroform and isopropanol were doubled to account for the increased sample volume and pellets were resuspended in 30 μl of 10 mM Tris-HCl (pH 8.0). Samples were placed in a heat block set to 95°C for 2 min and then snap-cooled on ice for 1 min. 30 μl of 2X Equilibration Buffer (2X Transcription Buffer, 20 mM MgCl2) containing either no ligand or 500 μM ppGpp was mixed with each sample and incubated at 37°C for 20 min. Samples were then probed with BzCN as described in the section *RNA chemical probing*, purified by TRIzol LS extraction as described in the section *RNA purification*, and resuspended in 5 μl 10% (v/v) DMSO. All downstream sample processing steps were then followed exactly as described for TECprobe-ML experiments.

### Validation of dsDNA purification using homemade SPRI beads

To assess the effect of SPRI bead ratio on dsDNA recovery, increasing volumes of SPRI beads were mixed with 1 μl of a 4-fold dilution of 100 bp DNA ladder (New England Biolabs) and diluted to a final volume of 100 μl with 10 mM Tris-HCl (pH 8.0). Samples were incubated at room temperature for 5 min and purified as described in the section *SPRI bead purification of DNA*. DNA was eluted by resuspending beads in 10 μl of 10 mM Tris-HCl (pH 8.0), allowing the sample to sit undisturbed for 3 min, placing the tube on a magnetic stand for 1 min so that the beads collected on the tube wall, and transferring the supernatant into a microcentrifuge tube that contained 2 μl 6X DNA Loading Dye [10 mM Tris-HCl (pH 8.0), 30% (v/v) glycerol, 0.48% (w/v) SDS, 0.05% (w/v) Bromophenol Blue]. Samples were run on native TBE-polyacrylamide gels, stained with SYBR Gold Nucleic Acid Stain, and scanned using an Azure Biosystems Sapphire Biomolecular Imager as described previously^52^.

### Optimization of single-stranded nucleic acid purification using homemade SPRI beads

Single-stranded nucleic acid purification using SPRI beads was optimized by assessing the effect of isopropanol and PEG 8000 concentration on fragment size recovery. A mixture containing 1 μl of a 4-fold dilution of Low Range ssRNA Ladder (New England Biolabs) and 0.75 pmol each of RPIX_SC1_Bridge, dRP1_NoMod.R, and PRA1_shrt.F oligos (Supplementary Table 1) was combined with 22.5 μl SPRI beads, variable volumes of anhydrous isopropanol and 50% (w/v) PEG 8000, and RNase-free water to 100 μl.

Samples were incubated at room temperature for 5 min and purified as described in the section *SPRI bead purification of DNA* above, except the beads were washed with 80% (v/v) ethanol instead of 70% (v/v) ethanol. Nucleic acids were eluted by resuspending the beads in 5 μl of 10 mM Tris-HCl (pH 8.0), allowing the tube to sit undisturbed for 3 min, placing the tube on a magnetic stand for 1 min so that the beads collected on the tube wall, and transferring the supernatant into a microcentrifuge tube that contained 15 μl of formamide loading dye. Samples were then analyzed as described in the section *Denaturing Urea-PAGE*.

### Reverse transcription efficiency assay

Reverse transcription (RT) efficiency assays that tested the ability of betaine, formamide, and ET SSB to eliminate primer dimer and promote full length cDNA synthesis were performed as follows: a *Cbe pfl* ZTP riboswitch DNA template lacking the terminator poly-U tract and containing an internal biotin-TEG modification was transcribed and RNA was purified as described in the sections *Single-round in vitro transcription for TECprobe-ML and TECprobe-SL* and *RNA purification* using the TECprobe-SL procedure, except that transcription volumes were 25 μl and chemical probing was not performed. The dRP1_5Bio.R oligo was annealed to the 3’ end of the RNA and immobilized on equilibrated Streptavidin C1 beads as described in the section *cDNA synthesis and cleanup*. Reverse transcription was then performed using either SuperScript II or SuperScript III as described below.

All SuperScript II reactions contained 50 mM Tris-HCl (pH 8.0), 75 mM KCl, 0.5 mM dNTP Solution Mix, 10 mM DTT, 3 mM MnCl2, 0.1% (v/v) Triton X-100, and 5 U/μl SuperScript II. The beads were resuspended in in 14.5 μl of SuperScript II master mix, which omitted RT additives and SuperScript II, mixed with 5 μl of an additive solution (described below), and pre-heated at 42°C in a thermal cycler for 2 min before 0.5 μl of SuperScript II was added to the reaction.

All SuperScript III reactions contained 50 mM Tris-HCl (pH 8.0), 75 mM KCl, 0.5 mM dNTP Solution Mix, 5 mM DTT, 3 mM MnCl2, 0.1% (v/v) Triton X-100, and 5 U/μl SuperScript III. The beads were resuspended in 15 μl of SuperScript III master mix and then mixed with 5 μl of an RT additive solution.

When present, betaine was included at 1.25 M, formamide was included at 2% or 5% (v/v), and ET SSB was included at 10 ng/μl. All reverse transcription reactions were incubated at 42°C for 50 min, then at 70°C for 15 min to heat inactivate reverse transcriptase, and cooled to 12°C. Samples briefly spun down in a mini centrifuge, placed on a magnetic stand, and the supernatant was aspirated and discarded. The beads were resuspended in 25 μl Bead Elution Buffer [95% (v/v) formamide and 10 mM EDTA (pH 8.0)], heated at 100°C for 5 min, placed on a magnetic stand, and the supernatant was collected and added to 125 μl Stop Solution [0.6 M Tris-HCl (pH 8.0) and 12 mM EDTA (pH 8.0)]. Reactions were processed for denaturing urea-PAGE by phenol-chloroform extraction and ethanol precipitation. Briefly, 150 μl Phenol-Chloroform-Isoamyl Alcohol (25:24:1) (ThermoFisher) was added to the reaction and thoroughly vortexed. The sample was centrifuged at 18,500 x g and 4°C for 5 min and the aqueous phase was transferred to a new tube and precipitated by adding 15 μl 3 M Sodium Acetate (pH 5.5), 450 μl 100% Ethanol, and 1.5 μl Glycoblue Coprecipitant and stored at - 20°C overnight. The sample was centrifuged at 18,500 x g and 4°C for 30 min and the supernatant was aspirated and discarded. Pellets were resuspended with 15 μl formamide loading dye and analyzed as described in the section *Denaturing Urea-PAGE*.

### Intermediate fraction analysis

Visualization of RNA-to-cDNA processing was performed by collecting intermediate sample fractions for analysis by denaturing PAGE. For this analysis, individual samples were processed in parallel to maintain the sample volumes used in the final procedure. Briefly, terminally biotinylated *Cba* ppGpp template was transcribed in the absence of ligand and purified as described in the section *Transcription antitermination assays*. One sample was resuspended with 15 μl of formamide loading dye and the remaining samples were ligated to the 9N_VRA3 3’ adapter as described in the *RNA 3’ adapter ligation* section. Following ligation, one sample was TRIzol extracted and the pellet was resuspended with 15 of μl formamide loading dye. The remaining samples were purified as described in the section *SPRI bead purification of RNA*. One sample was TRIzol extracted after each SPRI bead purification (for a total of two samples extracted), and the pellets were resuspended in 15 μl of formamide loading dye. The remaining samples were annealed to dRP1_5Bio.R oligo and immobilized on equilibrated Streptavidin C1 beads as described in the section *cDNA synthesis and cleanup*. Samples were briefly spun down in a mini centrifuge and placed on a magnetic stand to dispose of the supernatant. The beads from one sample were resuspended in 25 μl Bead Elution Buffer, heated at 100°C for 5 min, placed on a magnetic stand and the supernatant was collected and added to 125 μl Stop Solution. The remaining samples were reverse transcribed and purified as described in the *cDNA synthesis and cleanup*, except that the beads from one sample were resuspended in 25 μl Bead Elution Buffer, instead of Storage Buffer, heated at 100°C for 5 min, placed on a magnetic stand and the supernatant was collected and added to 125 μl Stop Solution. The remaining reaction was washed with 75 μl of Storage Buffer, resuspended in 25 μl Storage Buffer, and stored at -20°C. Both samples in Stop Solution were phenol-chloroform extracted and ethanol precipitated, and the resulting pellets were each resuspended with 15 μl formamide loading dye. The samples resuspended in formamide loading dye were analyzed as described in the section *Denaturing Urea-PAGE*. The sample resuspended in Storage Buffer was processed and analyzed as described in the *Test amplification of TECprobe libraries* section below except that serial 4-fold dilutions of bead-bound cDNA libraries were amplified for only 21 cycles.

### Test amplification of TECprobe libraries

The number of PCR amplification cycles needed for TECprobe libraries was determined by performing a test amplification adapted from Mahat et al.^55^. Briefly, 14 μl of PCR master mix was added to 6 μl of a 16-fold dilution of the bead-bound cDNA libraries, such that the final concentration of components in the 20 μl PCR were: 1X Q5 Reaction Buffer (New England Biolabs), 1X Q5 High GC Enhancer (New England Biolabs), 200 μM dNTP Solution Mix, 250 nM RPIX Forward Primer (Supplementary Table 1) 250 nM dRP1_NoMod.R Reverse Primer (Supplementary Table 1), 10 nM RPIX_SC1_Bridge (Supplementary Table 1), and 0.02 U/μl Q5 High-Fidelity DNA Polymerase. Amplification was performed for 21 or 25 cycles, at an annealing temperature of 62°C and an extension time of 20 s. 20 μl of each supernatant was run on native TBE-polyacrylamide gels to assess both the fragment size distribution of the libraries and to determine the appropriate number of cycles for amplification of libraries for high-throughput sequencing. A representative gel image is shown in Supplementary Figure 3d.

### Preparation of dsDNA libraries for sequencing

Amplification of cDNA libraries for high throughput sequencing was performed by preparing separate 50 μl PCRs for each (+) and (-) sample that contained 1X Q5 Reaction Buffer, 1X Q5 High GC Enhancer, 200 μM dNTP Solution Mix, 250 nM RPI Indexing Primer (Supplementary Table 1), 250 nM dRP1_NoMod.R Reverse Primer (Supplementary Table 1), 10 nM SC1Brdg_MINUS or SC1Brdg_PLUS channel barcode oligo (Supplementary Table 1), 12 μl of bead-bound cDNA library, and 0.02 U/μl Q5 High-Fidelity DNA Polymerase. Amplification was performed as indicated above, using the number of cycles determined by the test amplification. Supernatants from completed PCRs were each mixed with 100 μl of SPRI beads and purified as described in *SPRI bead purification of DNA*. DNA was eluted into 20 μl of 10 mM Tris-HCl (pH 8.0), mixed with 40 μl of SPRI beads, and purified as described in *SPRI bead purification of DNA* a second time. Twice-purified DNA was eluted into 10 μl of 10 mM Tris-HCl (pH 8.0) and quantified using the Qubit dsDNA HS Assay Kit (Invitrogen) with a Qubit 4 Fluorometer. Molarity was estimated using the length distribution observed during test amplification.

### High-throughput DNA sequencing

Sequencing of chemically probed libraries was performed by Novogene Co. on an Illumina HiSeq 4000 System using 2x150 PE reads with 10% PhiX spike in. TECprobe-ML libraries were sequenced at a depth of ∼30 to ∼60 million PE reads. TECprobe-SL libraries were sequenced at a depth of ∼1 to ∼2 million PE reads.

### Sequencing read pre-processing using cotrans_preprocessor

After comparing replicate data sets that were analyzed individually (Figure 2e-g and Supplementary Figure 4d- f), TECprobe-ML replicate data were concatenated and analyzed together. cotrans_preprocessor handles target generation, sequencing read pre-processing, and ShapeMapper2 run script generation for TECprobe- ML and TECprobe-SL data analysis. Source code and documentation for cotrans_preprocessor are available at https://github.com/e-strobel-lab/TECtools/releases/tag/v1.0.0.

### Target generation for TECprobe-ML and TECprobe-SL experiments using cotrans_preprocessor

For TECprobe-ML experiments, two types of sequence targets (3’ end targets and intermediate transcript targets) were generated by running cotrans_preprocessor in `MAKE_3pEND_TARGETS` mode. 3’ end targets files contained the 3’-most 14 nt of every intermediate transcript and all 1 nt substitution, insertion, and deletion variants of these sequences. The 3’ end targets file is used to demultiplex fastq files based on intermediate transcript identity, which is inferred from the RNA 3’ end, in the section *Sequencing read preprocessing for TECprobe-ML experiments*, below. The accuracy of 3’ end mapping was assessed by generating native and randomized variant test data using the options `-T` or `-T -R -U 30`, respectively, and processing the test data as described below in the section *Sequencing read preprocessing for TECprobe-ML experiments*. Based on these analyses, the 14 nt default length of the 3’-end target sequences was sufficient for accurate intermediate transcript identity determination. Intermediate transcript targets comprise individual fasta files for every intermediate transcript sequence with an A+1T+2 dinucleotide and the SC1 adapter appended to the 5’ end in lower case, so that these sequences are excluded from ShapeMapper2 analysis. For TECprobe-SL experiments, a single target was generated by running cotrans_preprocessor in MAKE_SINGLE_TARGET mode.

### Sequencing read pre-processing for TECprobe-ML experiments

For TECprobe-ML experiments, cotrans_preprocessor manages adapter trimming using fastp^56^ and demultiplexes aggregate sequencing reads by modified and untreated channel and by intermediate transcript identity. It was necessary to demultiplex aggregate sequencing reads by intermediate transcript identity, which is inferred from the RNA 3’ end, so that each intermediate transcript could be analyzed separately by ShapeMapper2; this avoids sequencing read multi-mapping during bowtie2 alignment. TECprobe-ML sequencing reads were processed by running cotrans_preprocessor in `PROCESS_MULTI` mode, which performs the following operations: First, fastp is called to perform adapter trimming and to extract unique molecular index and channel barcode sequences from the head of reads 1 and 2, respectively. After fastp processing is complete, cotrans_preprocessor splits the fastp output files by channel (modified barcode [RRRYY] or untreated barcode [YYYRR]) and intermediate transcript identity, and generates the files `smooth_transition.sh` and `config.txt`. `smooth_transition.sh` is a shell script that can be used to apply neighboring transcript smoothing by generating fastq files in which, for each intermediate transcript *n*, sequencing reads for intermediate transcripts *n-1*, *n*, and *n+1* are concatenated into a single file (excluding the minimum transcript *min*, which is concatenated as *min*, *min+1*, and the maximum transcript *max*, concatenated as *max-1*, *max*). `config.txt` is a configuration file that is used to specify information about a data set when generating a ShapeMapper2^57^ run script as described below in the section *ShapeMapper2 run script generation using cotrans_preprocessor*.

### Sequencing read pre-processing for TECprobe-SL experiments

For TECprobe-SL experiments, cotrans_preprocessor manages adapter trimming using fastp^56^ and demultiplexes aggregate sequencing reads by modified and untreated channels. TECprobe-SL sequencing reads were processed by running cotrans_preprocessor in `PROCESS_SINGLE` mode, which performs the following operations: First, fastp is called to perform adapter trimming and to extract the channel barcode sequence from the head of read 2. After fastp processing is complete, cotrans_preprocessor splits the fastp output files by channel (modified barcode [RRRYY] or untreated barcode [YYYRR]) and generates the file `config.txt`. As described above in the section *Sequencing read pre-processing for TECprobe-ML experiments*, `config.txt` is a configuration file that is used to specify information about a data set when generating a ShapeMapper2^57^ run script.

### ShapeMapper2 run script generation using cotrans_preprocessor

Shell scripts to run ShapeMapper2^57^ analysis were generated by running cotrans_preprocessor in `MAKE_RUN_SCRIPT` mode. For TECprobe-SL experiments, the run script contained a single command. For TECprobe-ML experiments, the run script contained a command for every intermediate transcript. The following ShapeMapper2 options were used: `--min-depth 500` was used to ensure that all data exceeded the minimum cutoff so that sequencing depth filtering could be performed separately during heatmap generation. `--min-qual-to-trim 10` was used to keep read pairs in which read 2 contained low quality base calls near the read head. `--min-qual-to-count 25` was used to filter low quality base calls during reactivity calculation.

### TECprobe-ML reactivity heatmap generation

ShapeMapper2 output data for each individual transcript in a TECprobe-ML experiment were assembled into a single matrix in csv format by the `compile_SM2_output` script. Reactivity and read depth data matrices and heatmaps were then generated by the `generate_cotrans_heatmap` script. Some transcript lengths were not enriched because the template DNA strand segment composed of the HP4_5bio.R primer does not contain randomly positioned internal biotin modifications. These transcripts were excluded from the heatmaps and all analyses. Source code and documentation for `compile_SM2_output` and `generate_cotrans_heatmap` are available at https://github.com/e-strobel-lab/TECprobe_visualization.

### Generation of correlation plots for TECprobe-ML data sets

Reactivity matrices generated by the `generate_cotrans_heatmap` script were used to generate replicate correlation plots using the `plot_cotrans_correlation` script. Transcript lengths that were not enriched by biotin-streptavidin roadblocking (as described above in *TECprobe-ML reactivity heatmap generation*) were excluded from the analysis. Nucleotides at RNA 3’ ends with a reactivity value of zero were excluded from the analysis. If the reactivity of a nucleotide in one replicate was NaN, the corresponding reactivity in the other replicate was masked as NaN. Source code and documentation for `plot_cotrans_correlation` are available at https://github.com/e-strobel-lab/TECprobe_visualization.

### Comparison of reactivity and background mutation rates for neighboring transcript lengths

The correlation of reactivity values and background mutation rates for neighboring transcripts was determined using the `compare_cotrans_neighbors` script. Pearson’s correlation coefficients for reactivity and background mutation rate were calculated for each pair of neighboring transcripts; the 3’-most nucleotide of the longer transcript, which is not present in the shorter transcript, was omitted from the analysis. Transcript lengths that were not enriched by biotin-streptavidin roadblocking (as described above in *TECprobe-ML reactivity heatmap generation*) were excluded from the analysis. The set of correlation coefficients for each individual TECprobe-ML experiment were visualized using violin plots. Source code and documentation for `compare_cotrans_neighbors` are available at https://github.com/e-strobel-lab/TECprobe_visualization.

### Visualization of RNA secondary and tertiary structures

RNA secondary structure predictions were performed using the RNAstructure^58^ Fold command with default settings. Crystal structures were visualized using UCSF Chimera^59^.

## Data availability

Raw sequencing data that support the findings of this study have been deposited in the Sequencing Read Archive (https://www.ncbi.nlm.nih.gov/sra) with the BioProject accession code PRJNA929456. Individual BioSample accession codes are available in Supplementary Table 4. Fully processed reactivity data have been deposited in the RNA Mapping Database^60^ (https://rmdb.stanford.edu/). Individual accession codes for each data set are available in Supplementary Table 5. ShapeMapper2 output data have been deposited in Zenodo (<u>DOI: 10.5281/zenodo.7640593</u>). All other data that support the findings of this paper are available from the corresponding authors upon request.

## Code availability

TECtools can be accessed at https://github.com/e-strobel-lab/TECtools/releases/tag/v1.0.0. Scripts used for data visualization can be accessed at https://github.com/e-strobel-lab/TECprobe_visualization.

## Supporting information

This article contains supporting information.

## Author Contributions

E.J.S., conceptualization; C.E.S. and E.J.S., methodology; C.E.S., investigation; C.E.S. and E.J.S, Validation; C.E.S. and E.J.S, Formal Analysis; C.E.S. and E.J.S, Software; C.E.S. and E.J.S, writing – original draft; C.E.S. and E.J.S, writing – review & editing; E.J.S., supervision; E.J.S., funding acquisition.

## Materials Included

**Supplementary Figure 1.**
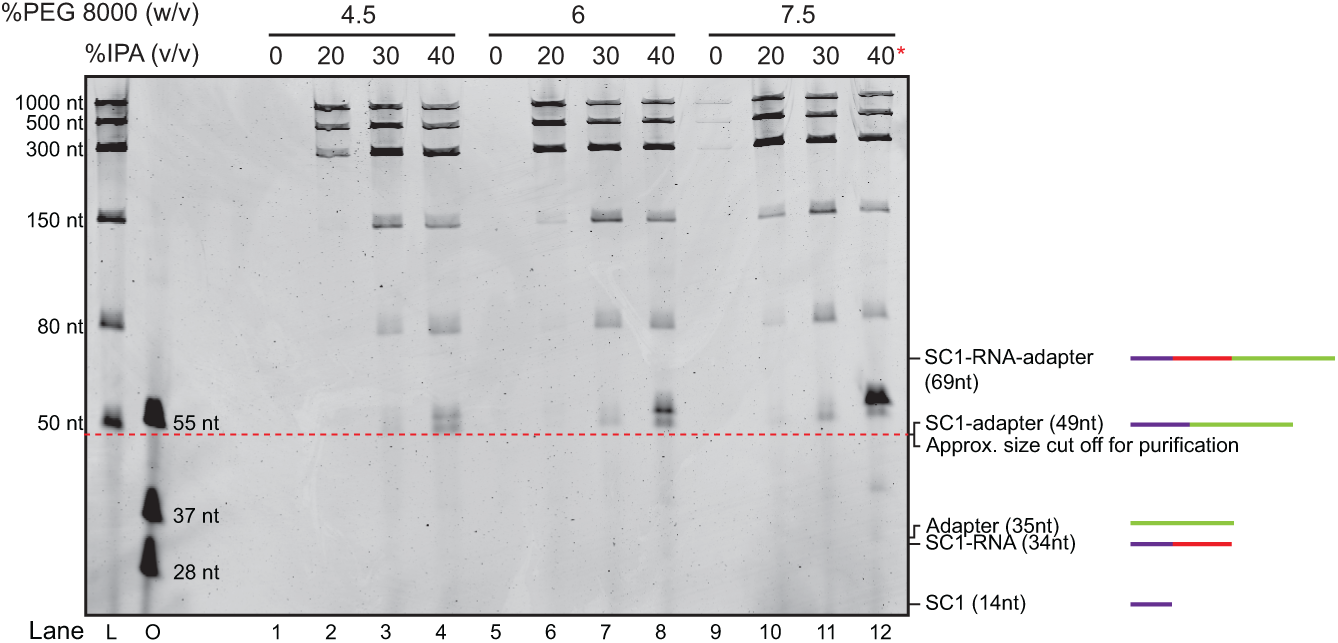
Purification of single stranded nucleic acids using SPRI beads. Denaturing PAGE analysis of nucleic acids purified using SPRI beads. Lanes L and O contain the Low Range ssRNA Ladder (New England Biolabs) and ssDNA oligonucleotide standards, respectively. The red asterisk marks the lane with the conditions used to purify RNA after the 3’ adapter ligation in TECprobe-ML experiments. Size diagrams of some RNAs of interest present in TECprobe-ML experiments are shown on the right. The 5’ SC1 hairpin (14nt - purple) is the smallest RNA before target RNA nucleotides are transcribed by RNAP. The first 19 target RNA transcript lengths are not analyzed in cotranscriptional structure probing experiments. Therefore, the SC1-RNA (34nt – purple and red) is the shortest RNA of interest and is a similar size to the adapter (35nt – green) before the RNA 3’ adapter ligation. Using the conditions in lane 12, the size cut off for fragment retention is approximately 50nt, which is similar to the combined length of the ligated 5’ SC1 hairpin and 3’ adapter (49nt – purple and green). The selected conditions retain most nucleic acids >50nt, which includes the smallest TECprobe-ML ligation product of interest (SC1-RNA-adapter 69nt – purple, red, and green) and deplete most smaller fragments. PEG, polyethylene glycol; IPA, isopropyl alcohol; SC1, structure cassette 1.

**Supplementary Figure 2.**
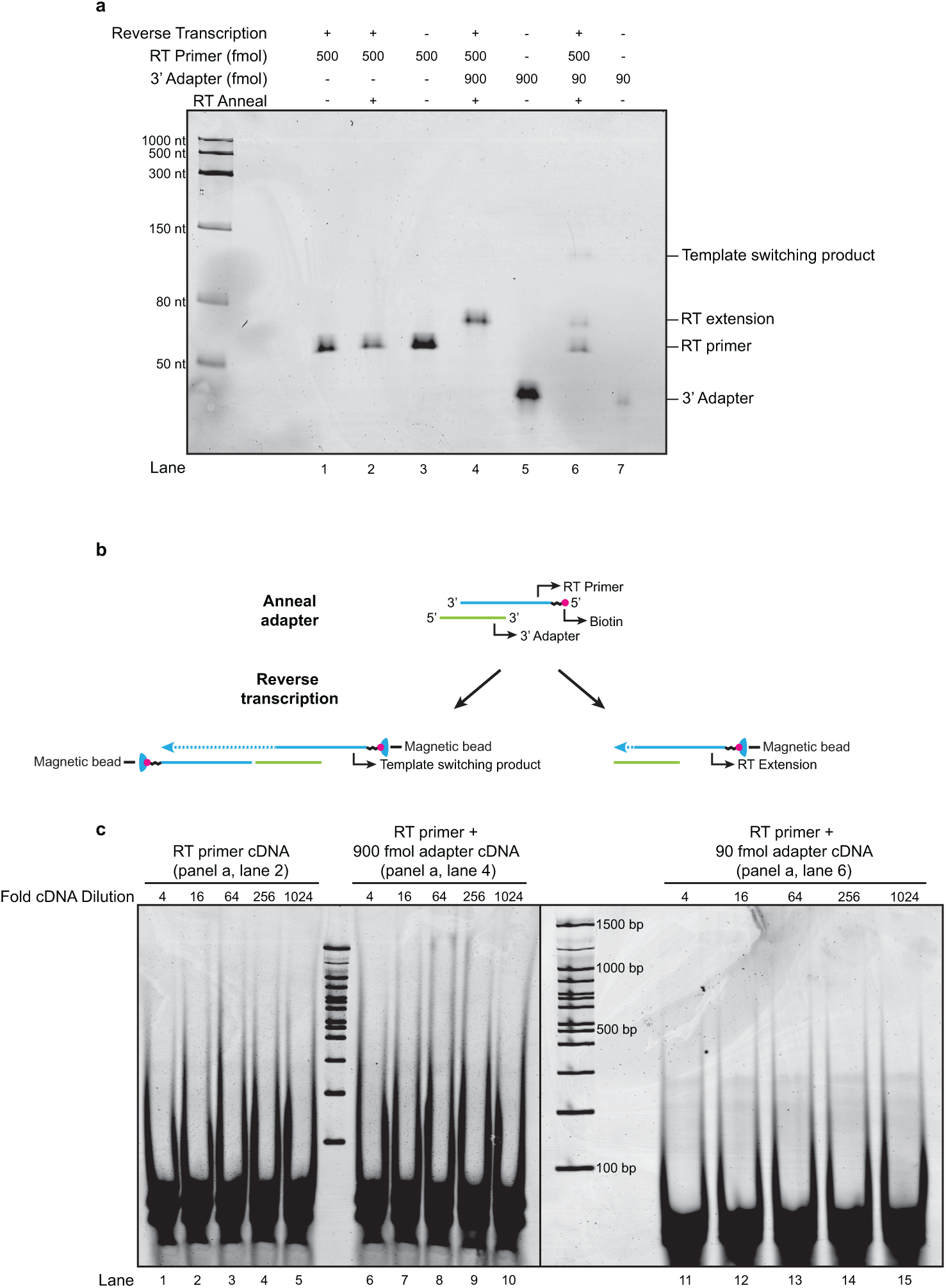
Characterization of reverse transcription primer dimer products that result from template switching. **a**, Denaturing PAGE analysis of reverse transcription reactions containing RT primer in the presence or absence of 3’ adapter. Lanes 3, 5 and 7 contain input nucleic acids for size comparison and were not subjected to reverse transcription. ‘RT anneal’ indicates whether the RT anneal protocol was used to denature nucleic acids and anneal the RT primer before reverse transcription. **b**, Illustration of the RT primer template switching and extension products. **c**, Native PAGE analysis of PCR amplification products using cDNA from lanes 2, 4, and 6 in Supplementary Figure 2a as templates. Samples were diluted as indicated and subjected to 21 cycles of PCR amplification. RT, reverse transcription.

**Supplementary Figure 3.**
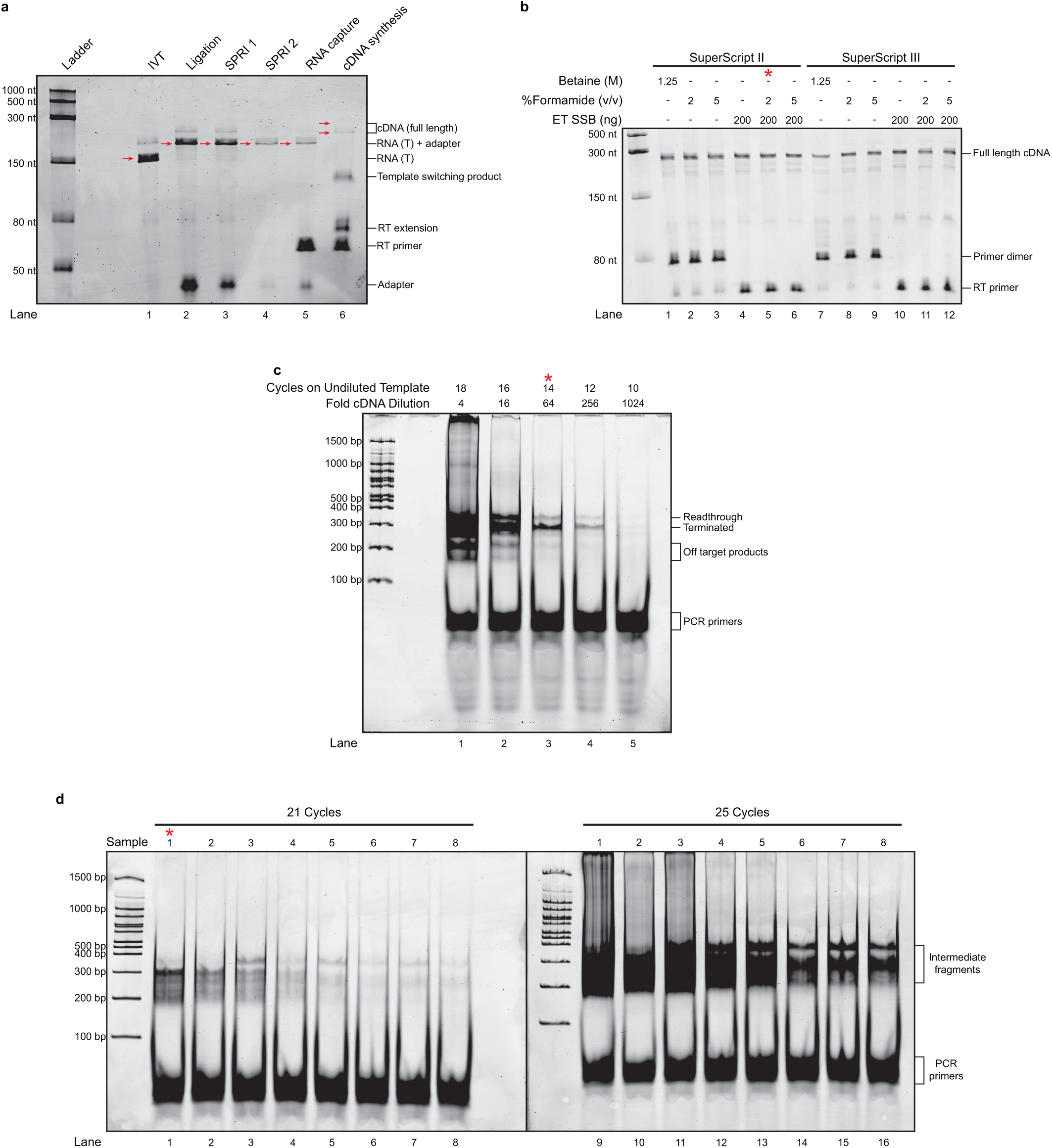
TECprobe RNA processing and cDNA amplification. **a**, Denaturing PAGE analysis of TECprobe-ML processing intermediates. *Cba* ppGpp riboswitch DNA templates without randomly biotinylated nucleotides were used to produce two prominent bands for easier visualization of sample processing. The identities of bands indicated by red arrows are shown on the right. **b**, Denaturing PAGE analysis of reverse transcription reactions using SuperScript II or III reverse transcriptase and various combinations of additives. A *pfl* ZTP riboswitch template containing a biotin-TEG stall site and inactivated transcription terminator was used. The red asterisk represents the conditions used for TECprobe experiments. **c**, Test amplification (21 cycles) using serial dilutions of the cDNA from panel a, lane 6. Readthrough and terminated products represent the PCR products of cDNA that was produced from readthrough and terminated transcripts, respectively. Fold cDNA dilution indicates the dilution of the cDNA, relative to the undiluted cDNA, that was subjected to PCR. Cycles on undiluted template indicates the number of cycles needed to achieve the same concentration of PCR product using 12 μl of undiluted cDNA. The red asterisk represents the lane with the appropriate band intensity to target when amplifying cDNA from TECprobe-SL experiments. **d**, Native PAGE analysis of dsDNA libraries generated using 1:16 diluted cDNA from TECprobe-ML experiments targeting the *Cba* ppGpp riboswitch after 21 cycles (left) or 25 cycles (right) of PCR. The red asterisk indicates the appropriate band intensity to target when ampliflying cDNA from TECprobe-ML experiments. IVT, *in vitro* transcription; SPRI, solid-phase reversible immobilization; T, terminated; RT, reverse transcription; ET SSB, extreme thermostable single-stranded DNA binding protein; TEG, triethylene glycol.

**Supplementary Figure 4.**
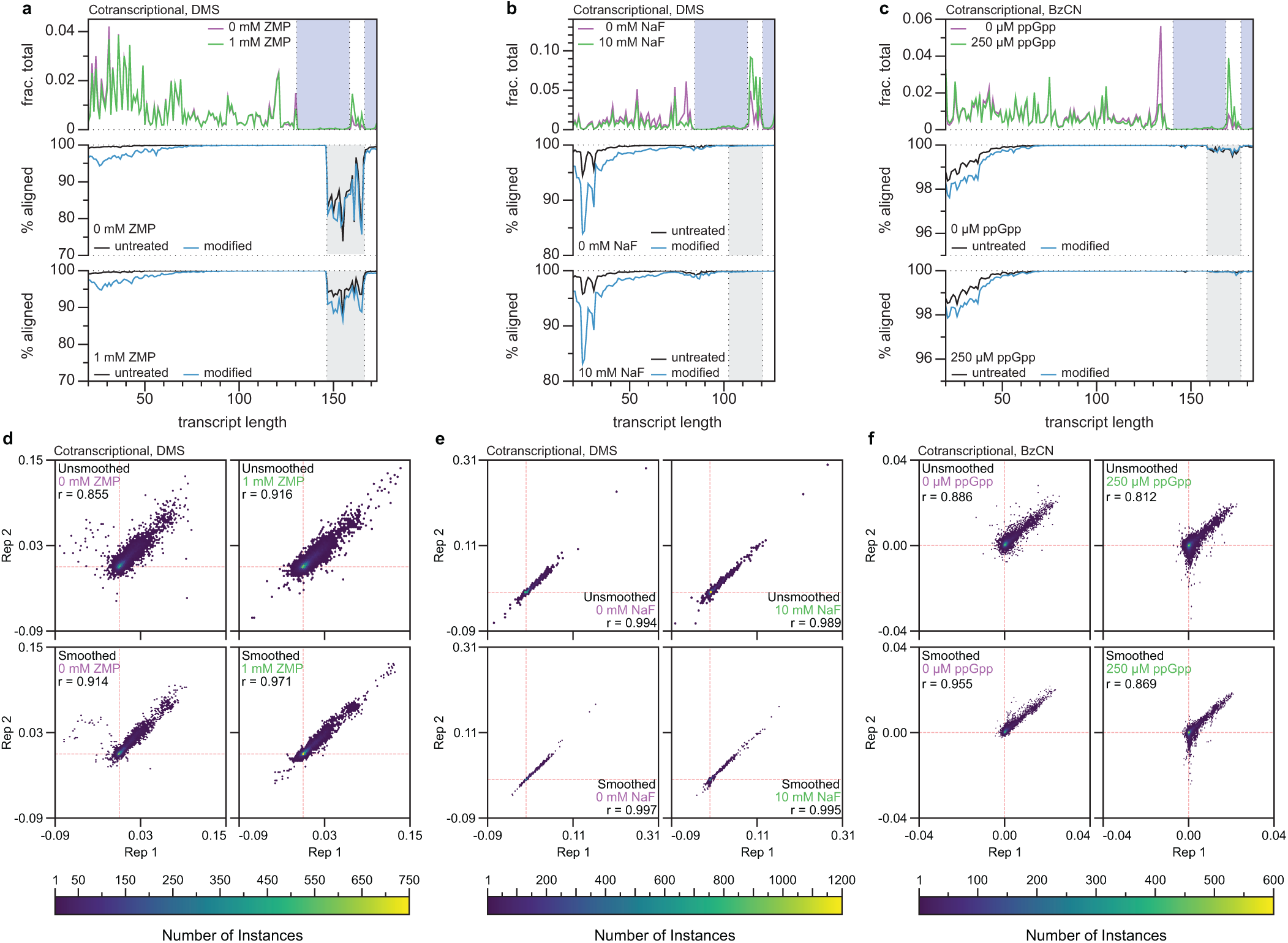
TECprobe-ML performance benchmarks. a-c, Plots showing the fraction of aligned reads that mapped to each transcript length (top) and the percentage of the reads kept after splitting those that aligned to each transcript length (middle, bottom) for the *Cbe pfl* ZTP riboswitch (DMS) (**a**), the *Bce crcB* fluoride riboswitch (DMS) (**b**), and the *Cba* ppGpp riboswitch (BzCN) (**c**) datasets. The blue shading in the top plot indicates transcripts that were not enriched by biotin-streptavidin roadblocking. The grey shading in the middle and lower plots indicates transcripts in which alignment was lower due to the presence of a nucleic acid species that is kept during fastq splitting but does not align. **d-f**, Hexbin plots comparing the reactivity of replicates for the *Cbe pfl* ZTP (DMS) (**d**), *Bce crcB* fluoride (DMS) (**e**), and *Cba* ppGpp (BzCN) (**f**) riboswitches for both unsmoothed (top) and smoothed (bottom) datasets. Data were plotted with a grid size of 75 by 75 hexagons, and the depth of overlapping data points are indicated by the heatmap. Red dashed lines indicate the position of 0 for each axis. BzCN, benzoyl cyanide; DMS, dimethyl sulfate.

**Supplementary Figure 5.**
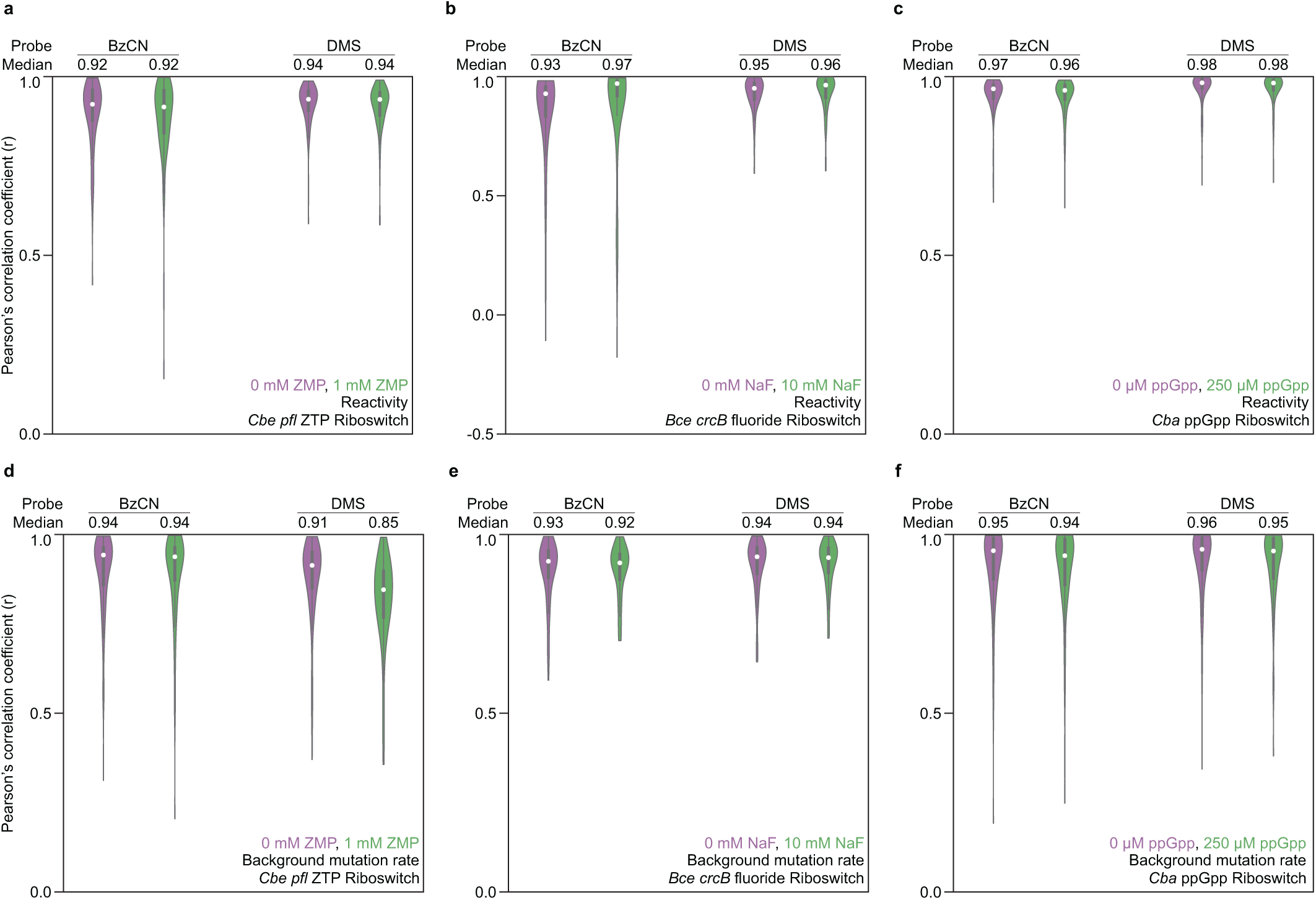
Comparison of reactivity and background mutation rates for neighboring transcripts. Violin plots of Pearson’s correlation coefficients comparing reactivities (**a-c**) or background mutation rates (**d-f**) of neighboring transcripts (*n* and *n*+1) for the *Cbe pfl* ZTP (**a** and **d**), the *Bce crcB* fluoride (**b** and **e**), and the *Cba* ppGpp (**c** and **f**) riboswitches. White circles indicate the data median. BzCN, benzoyl cyanide; DMS, dimethyl sulfate.

**Supplementary Figure 6.**
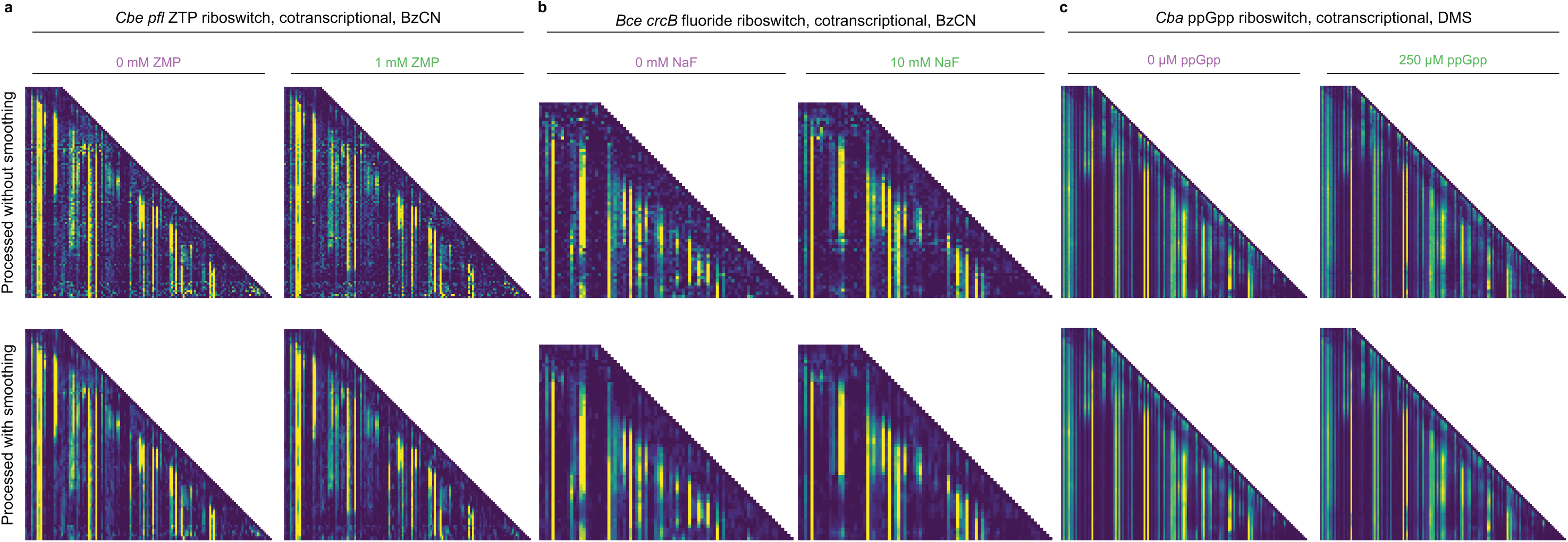
Comparison of reactivity matrices with and without neighboring transcript smoothing. TECprobe-ML reactivity matrices for the *Cbe pfl* ZTP (BzCN) (**a**), the *Bce crcB* fluoride (BzCN) (**b**), and the *Cba* ppGpp (DMS) (**c**) riboswitches. Matrices were generated from data processed without (top) or with (bottom) neighboring transcript smoothing. Reactivity is shown as background-subtracted mutation rate. BzCN, benzoyl cyanide; DMS, dimethyl sulfate.

**Supplementary Figure 7.**
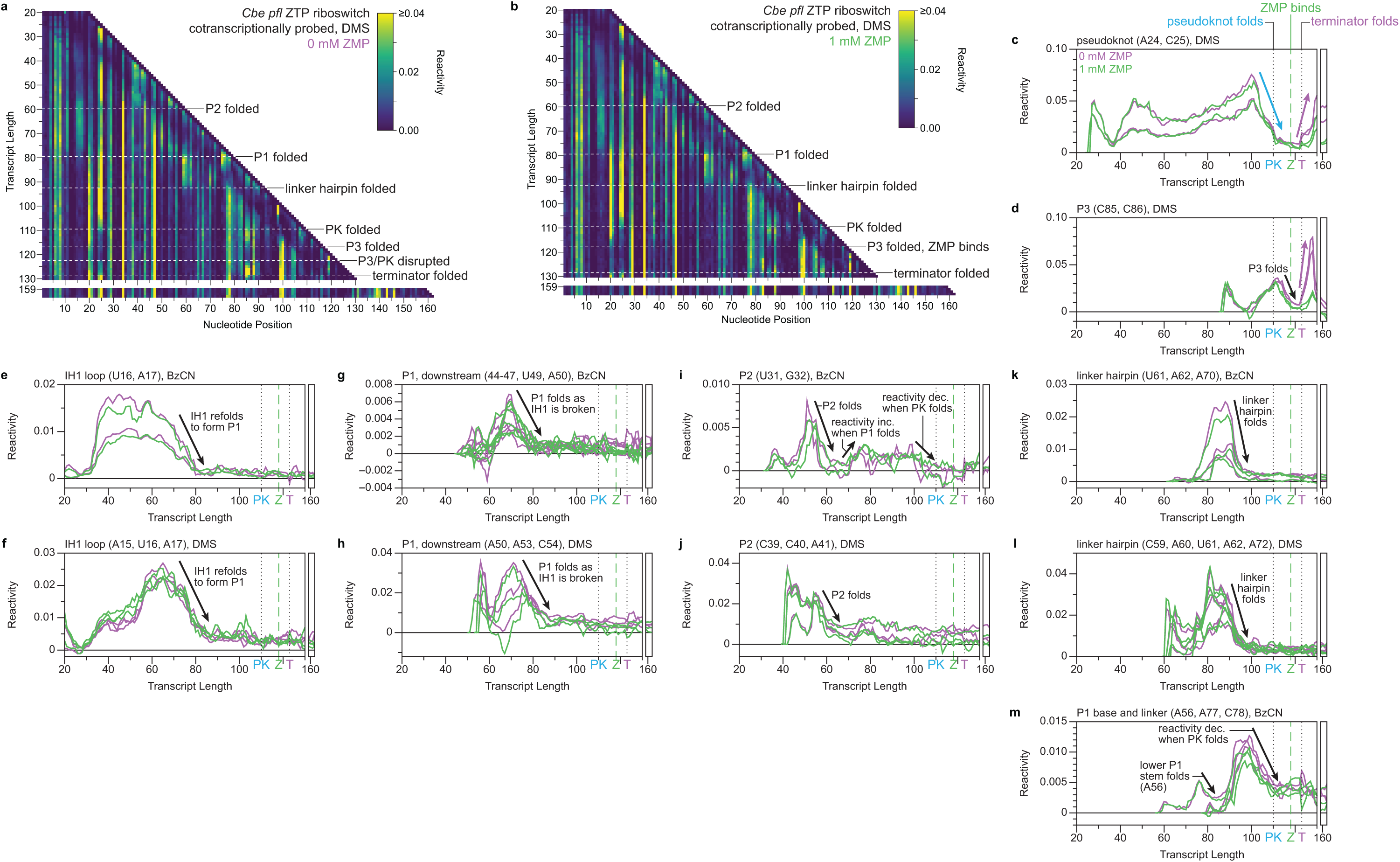
Cotranscriptional DMS probing of the *C*. *beijerinckii pfl* ZTP riboswitch and additional visualization of cotranscriptional folding transitions. **a**, **b**, TECprobe-ML DMS reactivity matrices for the *Cbe pfl* ZTP riboswitch with 0 mM and 1 mM ZMP. Transcripts 131-158, which were not enriched, are excluded. Reactivity is shown as background-subtracted mutation rate. Data are from two independent replicates that were concatenated and analyzed together. **c-m**, Transcript length-dependent reactivity changes showing folding transitions. BzCN data are from Figure 3c, d. Black dotted lines indicate pseudknot and terminator folding. Green dashed lines indicate ZMP binding. DMS, dimethyl sulfate; PK, pseudoknot; BzCN, benzoyl cyanide.

**Supplementary Figure 8.**
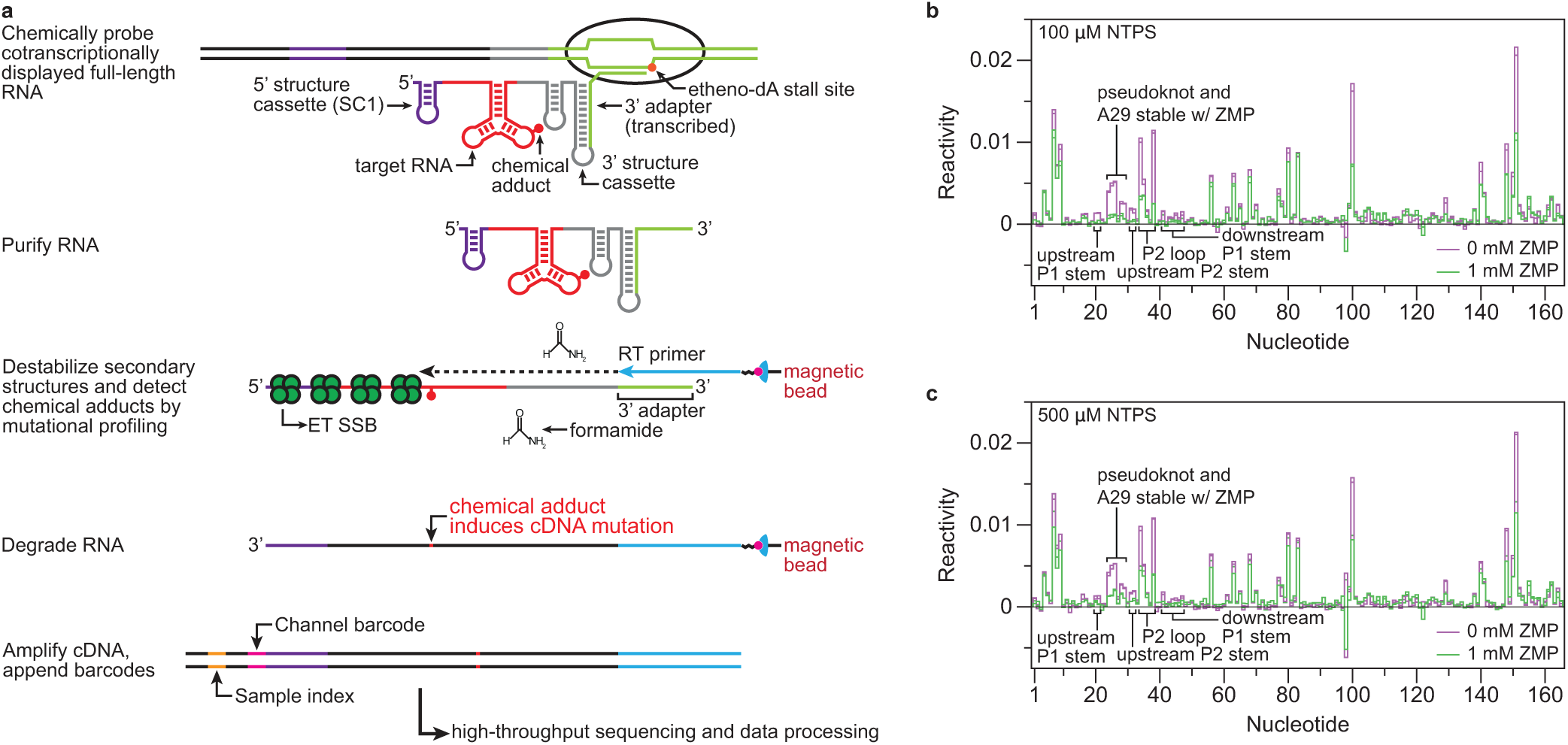
Cotranscriptional probing of full-length *C*. *beijerinckii pfl* ZTP riboswitch transcripts using TECprobe-SL. **a**, Illustration of the TECprobe-SL procedure. DNA templates that contain an internal etheno-dA modification are *in vitro* transcribed to generate roadblocked TECs that contain cotranscriptionally folded RNA. A 3’ structure cassette is used to sequester the 3’ adapter sequence from interactions that could influence target RNA structure. Cotranscriptionally displayed RNA is then chemically probed. The transcribed sequence contains the 3’ adapter so no ligation is needed. Following cotranscriptional chemical probing, solid-phase error-prone reverse transcription is performed. Full-length cDNA is amplified using primers that anneal to the 3’ adapter and the 5’ structure cassette. The resulting libraries are sequenced and processed using fastp^56^, TECtools, and ShapeMapper2^57^. **b**,**c** TECprobe-SL reactivity profiles for cotranscriptionally BzCN-probed full length *Cbe pfl* ZTP transcripts with 100 μM (**b**) or 500 μM (**c**) NTPs. The transcription terminator was inactivated by mutations to the poly-U tract to promote full-length RNA synthesis in the absence of ZMP. Two independent replicates are plotted for each condition. RT, reverse transcription; ET SSB, extremely thermostable single-stranded binding protein.

**Supplementary Figure 9.**
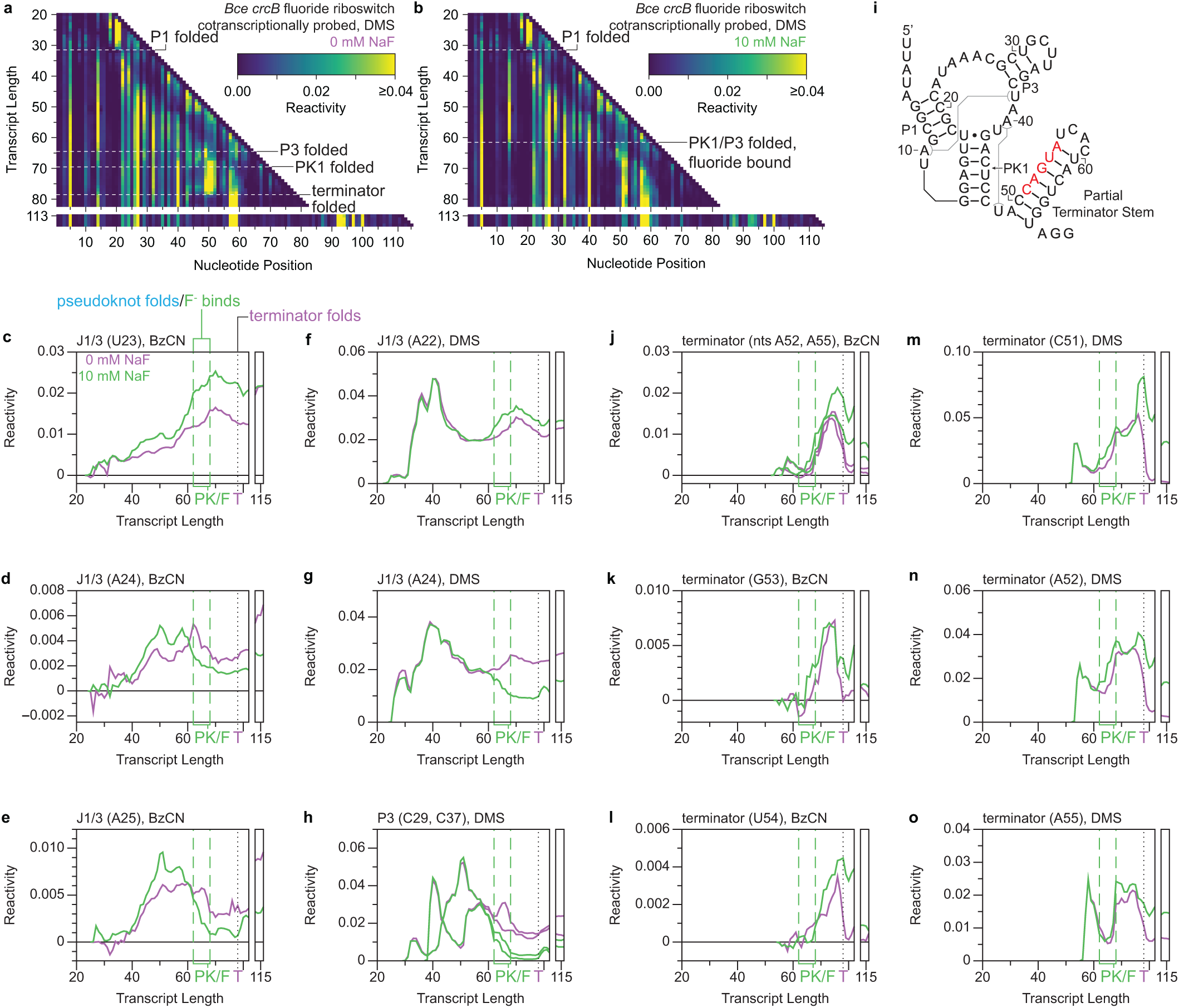
Cotranscriptional DMS probing of the *B. cereus crcB* fluoride riboswitch and additional visualization of cotranscriptional folding transitions. **a**,**b**, TECprobe-ML DMS reactivity matrices for the *Bce crcB* fluoride riboswitch with 0 mM and 10 mM NaF. Transcripts 83-112, which were not enriched, are excluded. Reactivity is shown as background-subtracted mutation rate. Data are from two independent replicates that were concatenated and analyzed together. **c-h**, Transcript length-dependent reactivity changes in J1/3 (**c-g**) and P3 (**h**). Green dashed lines indicate pseudoknot folding. The black dotted line indicates terminator folding. **i**, Secondary structure with partial nucleation of the terminator stem. Red nucleotides exhibit fluoride-dependent reactivity changes that are consistent with delayed terminator hairpin nucleation. **j-o**, Transcript length-dependent reactivity changes in the terminator. BzCN data are from Figure 4c, d. DMS, dimethyl sulfate; PK, pseudoknot; BzCN, benzoyl cyanide.

**Supplementary Figure 10.**
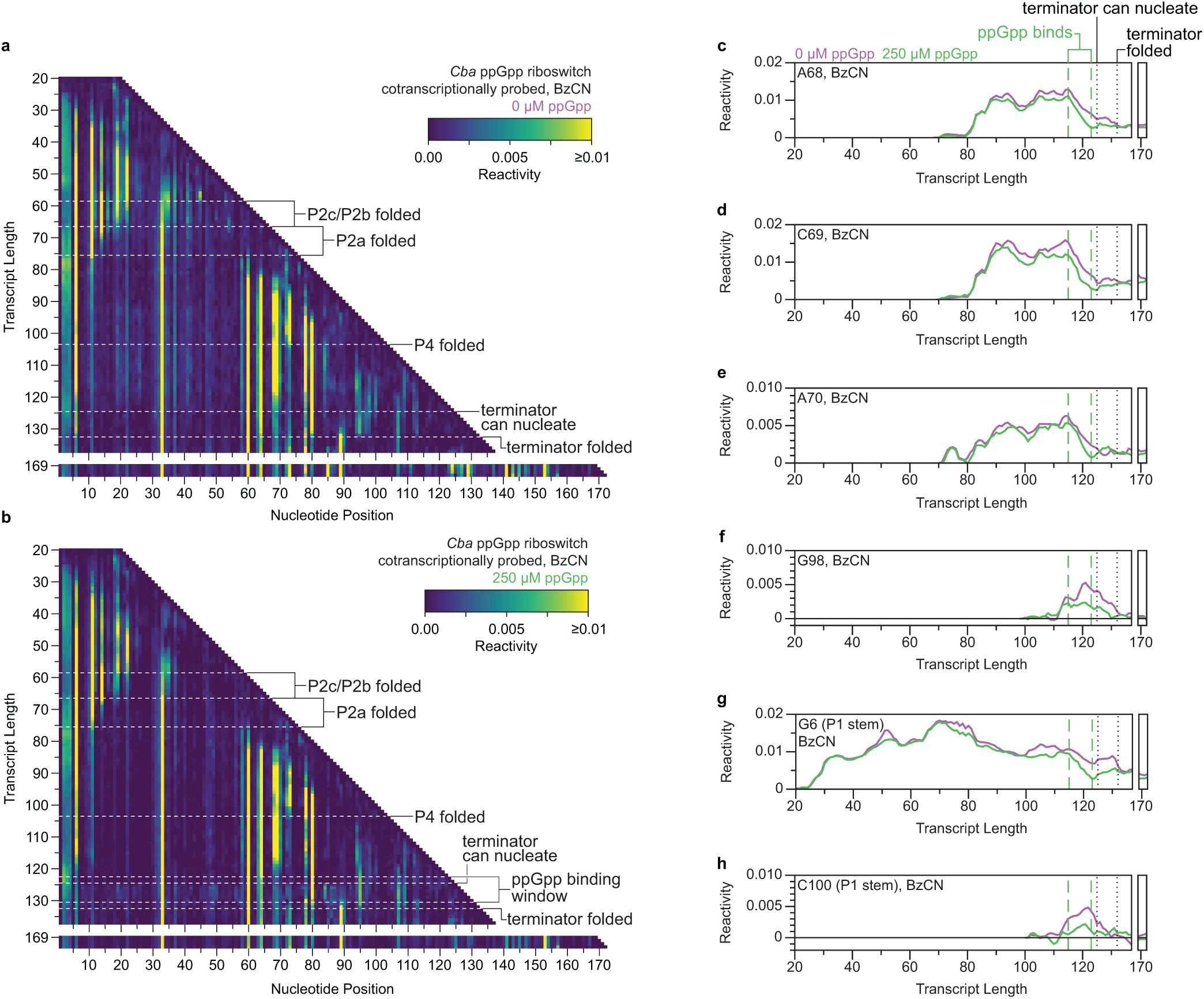
Cotranscriptional BzCN probing of the *C. bacterium* ppGpp riboswitch. **a**,**b**, TECprobe-ML BzCN reactivity matrices for the *Cba* ppGpp riboswitch with 0 μM and 250 μM ppGpp. Transcripts 138-168, which were not enriched, are excluded. The annotation ‘terminator can nucleate’ indicates the transcript at which a coordinated weak increase in reactivity is observed at numerous positions in the ppGpp aptamer when ppGpp is present, which can be seen in panel b. Reactivity is shown as background-subtracted mutation rate. Data are from two independent replicates that were concatenated and analyzed together. **c-h**, Transcript length-dependent reactivity changes in the ligand binding pocket (**c-e**), J1-4 (**f**), and P1 stem (**g, h**). Green dashed lines indicate the bounds of the apo-to-ppGpp-bound folding transition. Black dotted lines indicate positions where the terminator begins to nucleate and is folded. BzCN, benzoyl cyanide.

**Supplementary Figure 11.**
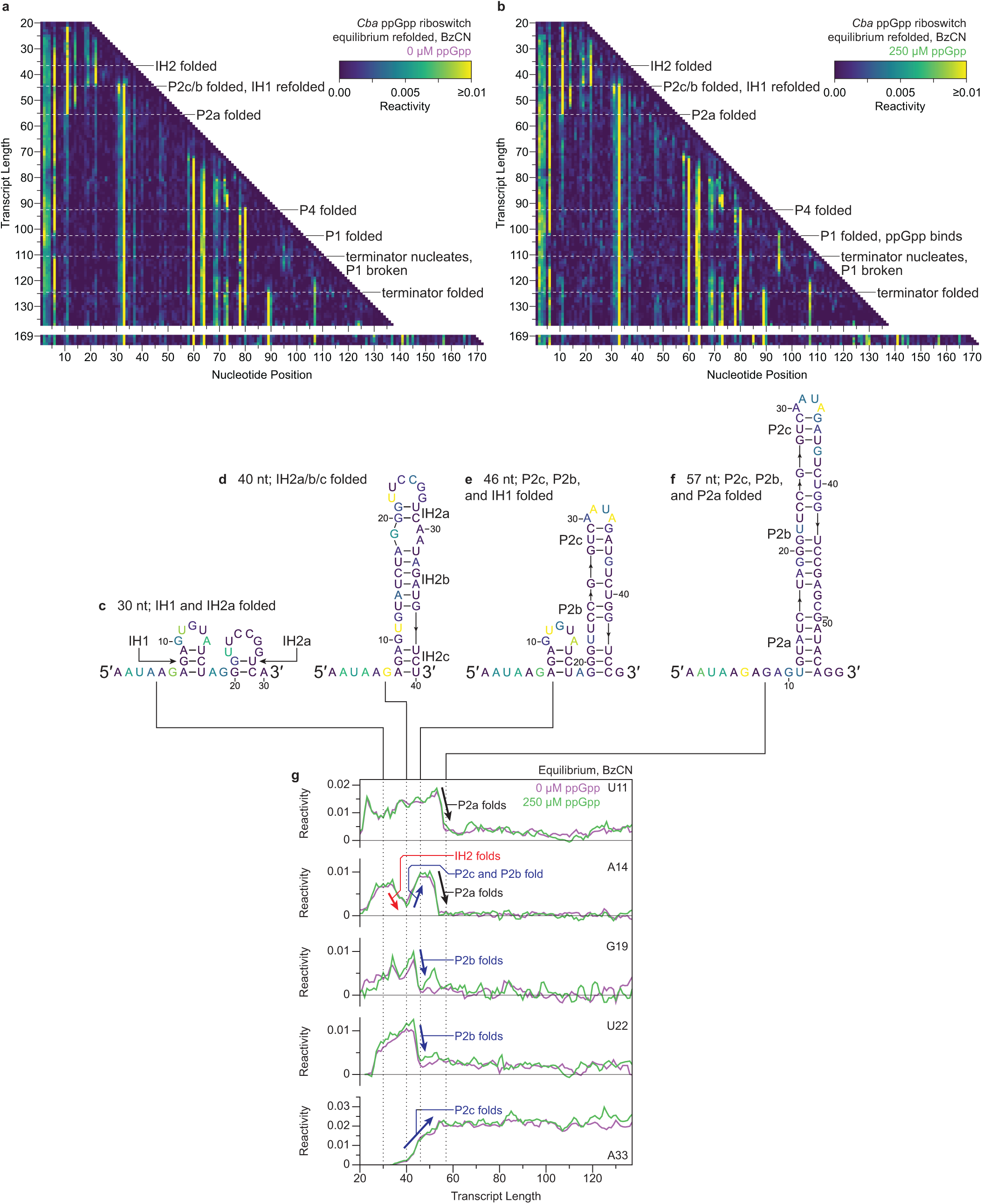
Equilibrium BzCN probing of the *C*. *bacterium* ppGpp riboswitch. **a**,**b**, TECprobe-ML BzCN reactivity matrices of *Cba* ppGpp riboswitch intermediate transcripts that were purified and refolded in the absence or presence of 250 μM ppGpp before chemical probing. Transcripts 138- 168, which were not enriched, are excluded. Reactivity is shown as background-subtracted mutation rate. (**c-f**) Secondary structures of proposed early folding intermediates colored by reactivity from transcripts 30 (**c**), 40 (**d**), 46 (**e**), and 57 (**f**) in the absence of ppGpp. **g**, Plots showing transcript length-dependent reactivity changes that occur during P2 hairpin folding. Black dotted lines indicated the transcript lengths used to identify the intermediate structures shown in **c-f**. BzCN, benzoyl cyanide.

**Supplementary Figure 12.**
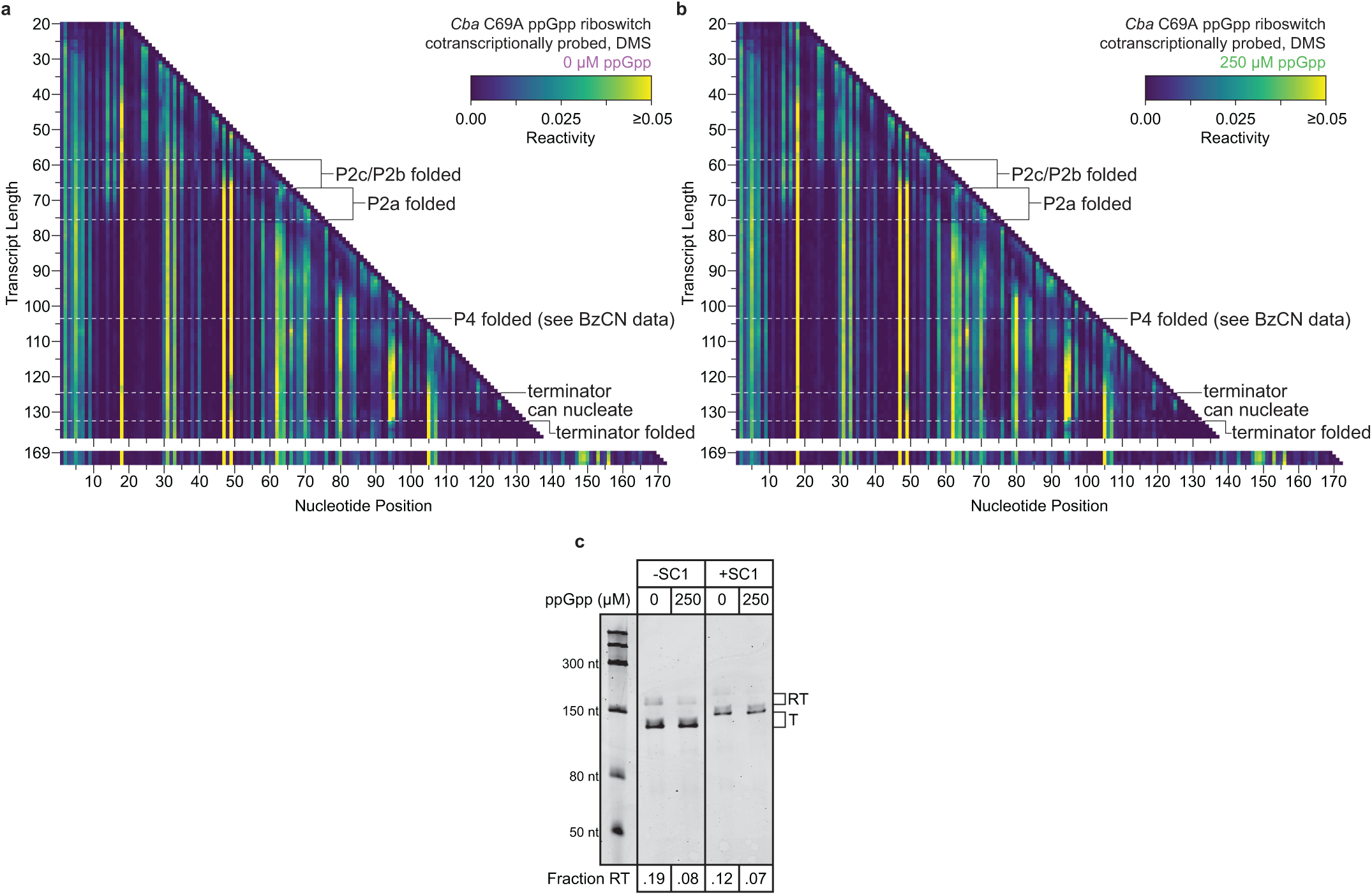
Cotranscriptional DMS probing of the *C. bacterium* ppGpp riboswitch C69A variant. **a**,**b**, TECprobe-ML DMS reactivity matrices for the *Cba* ppGpp riboswitch C69A variant with 0 μM and 250 μM ppGpp. Transcripts 138-168, which were not enriched, are excluded. Data are from two independent replicates that were concatenated and analyzed together. Reactivity is shown as background-subtracted mutation rate. **c**, Single-round transcription termination assays for the *Cba* ppGpp riboswitch C69A variant with or without SC1. Fraction readthrough is the average of two independent replicates. DMS, dimethyl sulfate; BzCN, benzoyl cyanide; SC1, structure cassette 1; RT, readthrough; T, terminated.

**Supplementary Figure 13.**
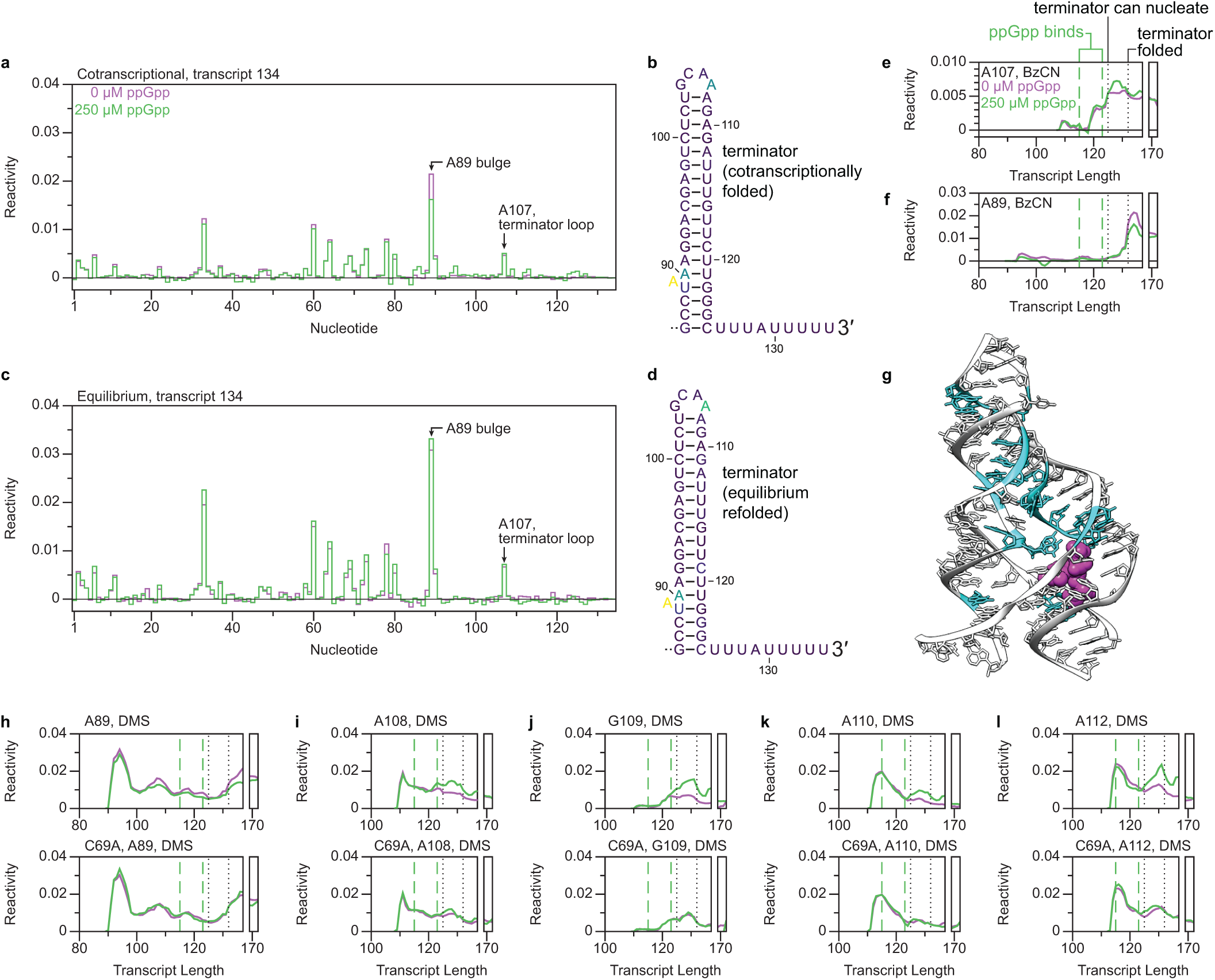
Visualization of *Cba* ppGpp terminator hairpin folding. a-d. , BzCN reactivity of *Cba* ppGpp riboswitch transcripts from the primary transcription termination site (nt 134) in cotranscriptional (**a**) and equilibrium (**c**) probing conditions. Terminator hairpin secondary structures are colored by the cotranscriptional (**b**) and equilibrium (**d**) reactivity of transcript 134. **e**,**f**, Transcript length-dependent BzCN reactivity changes that occur during cotranscriptional terminator folding. Green dashed lines indicate the bounds of the apo-to-ppGpp-bound folding transition. Black dotted lines indicate positions where the terminator begins to nucleate and is folded. Data are from Supplementary Figure 10a, b. **g**, Structure of the *Thermoanaerobacter mathranii* PRPP riboswitch G96A (ppGpp-binding) variant (PDB 6CK4)^38^ highlighted to indicate nucleotides that undergo a weak increase in BzCN reactivity at transcript 125 in the presence of ppGpp. **h-l**, Transcript length-dependent reactivity changes in terminator nucleotides. Data are from Figure 5d, e and Supplementary Figures 10a, 10b, 11a, 11b, 12a, and 12b. BzCN, benzoyl cyanide; DMS, dimethyl sulfate.

**Supplementary Table 1.**
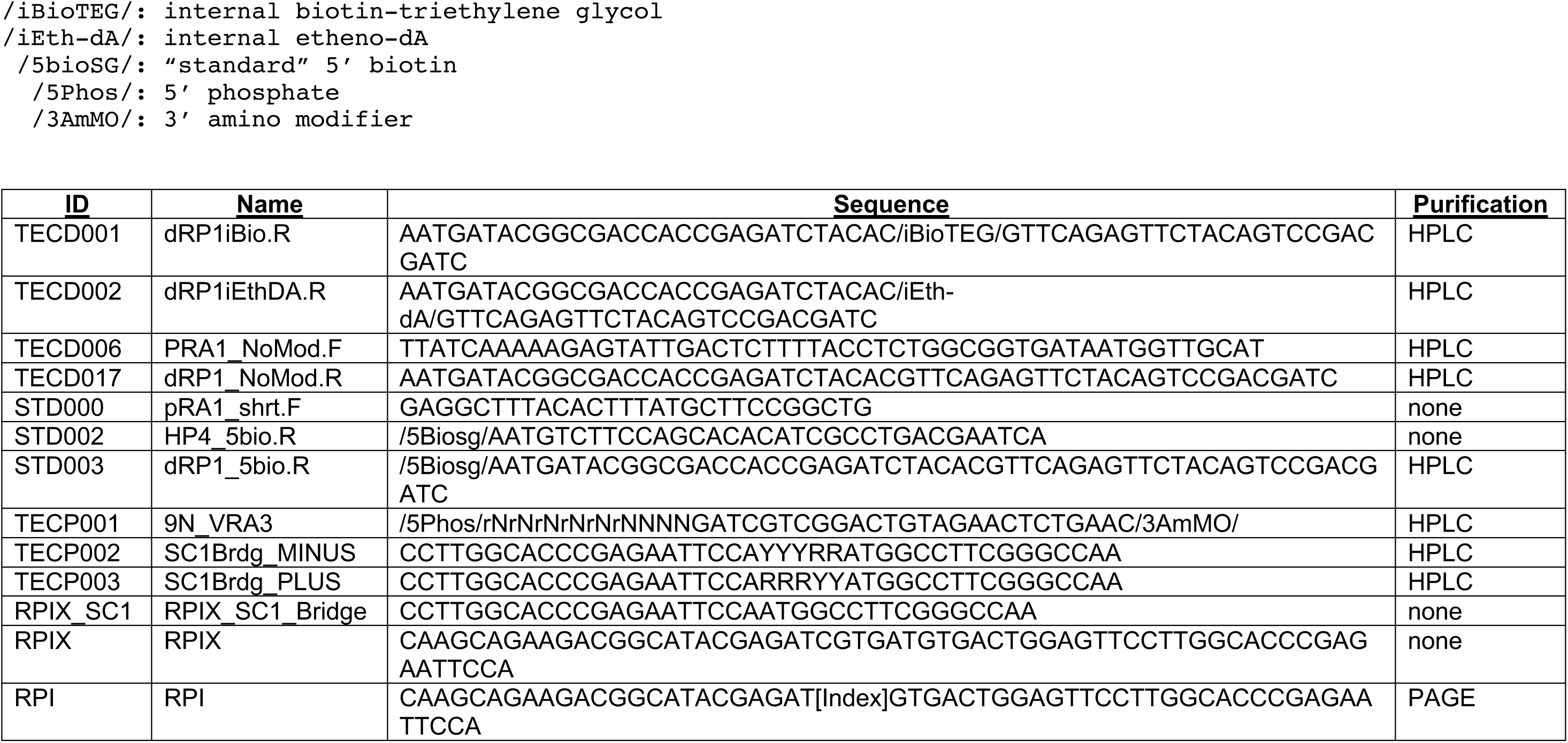
Oligonucleotides used in this study. The table below shows oligonucleotides used in this study. The modification codes presented are compatible with Integrated DNA Technology ordering.

**Supplementary Table S2.** DNA templates prepared for this study. The table below describes the DNA templates prepared for this study, including the primers and templates used, DNA modifications, whether translesion synthesis was performed, how the DNA template was purified, and the figures in which each DNA templates was used

| <u>ID</u> | <u>Fwd Primer</u> | <u>Rev Primer</u> | <u>Template</u> | <u>Modifications</u> | <u>Translesion Synthesis</u> | <u>Clean up</u> | <u>Used in Fig(s)</u> |
| --- | --- | --- | --- | --- | --- | --- | --- |
| 1 | TECD006 | STD002 | pCES007 | 5' biotin | N/A | Gel extracted | 3 |
| 2 | TECD006 | STD002 | pCES008 | 5' biotin | N/A | Gel extracted | 3 |
| 3 | TECD006 | STD002 | Template 2 | Internal biotin-11 nucleotides, 5' biotin | N/A | SPRI beads | 1, 2, 3, S4, S5, S6, S7 |
| 4 | TECD006 | STD002 | pCES009 | 5' biotin | N/A | Gel extracted | 4 |
| 5 | TECD006 | STD002 | pCES010 | 5' biotin | N/A | Gel extracted | 4 |
| 6 | TECD006 | STD002 | Template 5 | Internal biotin-11 nucleotides, 5' biotin | N/A | SPRI beads | 2, 4, S4, S5, S6, S9 |
| 7 | STD000 | STD002 | pCES011 | 5' biotin | N/A | Gel extracted | 5 |
| 8 | STD000 | STD002 | pCES012 | 5' biotin | N/A | Gel extracted | 5, S3 |
| 9 | STD000 | STD002 | Template 8 | Internal biotin-11 nucleotides, 5' biotin | N/A | SPRI beads | 2, 5, 6, 7, S3, S4, S5, S6, S10, S11, S13 |
| 10 | STD000 | STD002 | pCES013 | 5' biotin | N/A | Gel extracted | S12 |
| 11 | STD000 | STD002 | pCES014 | 5' biotin | N/A | Gel extracted | S12 |
| 12 | STD000 | STD002 | Template 11 | Internal biotin-11 nucleotides, 5' biotin | N/A | SPRI beads | 2, 7, S12, S13 |
| 13 | TECD006 | TECD002 | Gel-purified linear template from pCES015 | Internal etheno-dA | Yes | SPRI beads | S8 |
| 14 | TECD006 | TECD001 | Gel-purified linear template from pCES015 | Internal Bio-TEG | Yes | SPRI beads | S3 |

**Supplementary Table 3.**
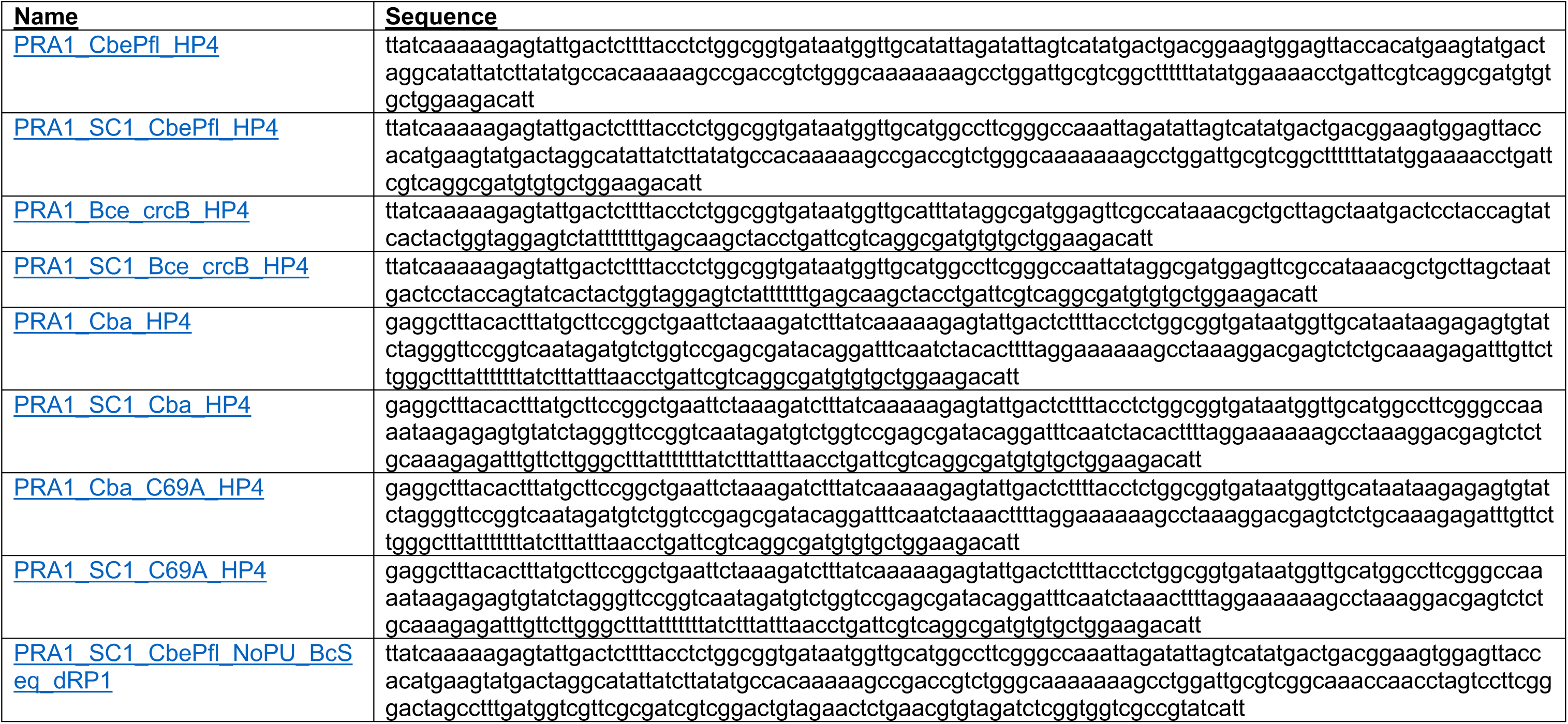
DNA template sequences. The table below contains the DNA sequences used in this study.

**Supplementary Table 4.** Sequencing Read Archive (SRA) deposition table. All primary sequencing data generated in this work are freely available from the Sequencing Read Archive (http://www.ncbi.nlm.nih.gov/sra), accessible via the BioProject accession number PRJNA929456 or using the individual accession numbers below.

| SRA Acession | Sample Name | Experiment |
| --- | --- | --- |
| SAMN32973014 | ZTP_BzCN_0mM_rep1 | wt ZTP riboswitch multilength cotranscriptional BzCN probing with 0mM ZMP, replicate 1 |
| SAMN32973015 | ZTP_BzCN_0mM_rep2 | wt ZTP riboswitch multilength cotranscriptional BzCN probing with 0mM ZMP, replicate 2 |
| SAMN32973016 | ZTP_BzCN_1mM_rep1 | wt ZTP riboswitch multilength cotranscriptional BzCN probing with 1mM ZMP, replicate 1 |
| SAMN32973017 | ZTP_BzCN_1mM_rep2 | wt ZTP riboswitch multilength cotranscriptional BzCN probing with 1mM ZMP, replicate 2 |
| SAMN32973018 | ZTP_DMS_0mM_rep1 | wt ZTP riboswitch multilength cotranscriptional DMS probing with 0mM ZMP, replicate 1 |
| SAMN32973019 | ZTP_DMS_0mM_rep2 | wt ZTP riboswitch multilength cotranscriptional DMS probing with 0mM ZMP, replicate 2 |
| SAMN32973020 | ZTP_DMS_1mM_rep1 | wt ZTP riboswitch multilength cotranscriptional DMS probing with 1mM ZMP, replicate 1 |
| SAMN32973021 | ZTP_DMS_1mM_rep2 | wt ZTP riboswitch multilength cotranscriptional DMS probing with 1mM ZMP, replicate 2 |
| SAMN32973022 | ZTP_SL_BzCN_0mM_lo_rep1 | wt ZTP riboswitch single length cotranscriptional BzCN probing with 0mM ZMP and 100 uM NTPs, replicate 1 |
| SAMN32973023 | ZTP_SL_BzCN_0mM_lo_rep2 | wt ZTP riboswitch single length cotranscriptional BzCN probing with 0mM ZMP and 100 uM NTPs, replicate 2 |
| SAMN32973024 | ZTP_SL_BzCN_0mM_hi_rep1 | wt ZTP riboswitch single length cotranscriptional BzCN probing with 0mM ZMP and 500 uM NTPs, replicate 1 |
| SAMN32973025 | ZTP_SL_BzCN_0mM_hi_rep2 | wt ZTP riboswitch single length cotranscriptional BzCN probing with 0mM ZMP and 500 uM NTPs, replicate 2 |
| SAMN32973026 | ZTP_SL_BzCN_1mM_lo_rep1 | wt ZTP riboswitch single length cotranscriptional BzCN probing with 1mM ZMP and 100 uM NTPs, replicate 1 |
| SAMN32973027 | ZTP_SL_BzCN_1mM_lo_rep2 | wt ZTP riboswitch single length cotranscriptional BzCN probing with 1mM ZMP and 100 uM NTPs, replicate 2 |
| SAMN32973028 | ZTP_SL_BzCN_1mM_hi_rep1 | wt ZTP riboswitch single length cotranscriptional BzCN probing with 1mM ZMP and 500 uM NTPs, replicate 1 |
| SAMN32973029 | ZTP_SL_BzCN_1mM_hi_rep2 | wt ZTP riboswitch single length cotranscriptional BzCN probing with 1mM ZMP and 500 uM NTPs, replicate 2 |
| SAMN32973030 | F_BzCN_00mM_rep1 | wt fluoride riboswitch multilength cotranscriptional BzCN probing with 0 mM NaF, replicate 1 |
| SAMN32973031 | F_BzCN_00mM_rep2 | wt fluoride riboswitch multilength cotranscriptional BzCN probing with 0 mM NaF, replicate 2 |
| SAMN32973032 | F_BzCN_10mM_rep1 | wt fluoride riboswitch multilength cotranscriptional BzCN probing with 10 mM NaF, replicate 1 |
| SAMN32973033 | F_BzCN_10mM_rep2 | wt fluoride riboswitch multilength cotranscriptional BzCN probing with 10 mM NaF, replicate 2 |
| SAMN32973034 | F_DMS_00mM_rep1 | wt fluoride riboswitch multilength cotranscriptional DMS probing with 0 mM NaF, replicate 1 |
| SAMN32973035 | F_DMS_00mM_rep2 | wt fluoride riboswitch multilength cotranscriptional DMS probing with 0 mM NaF, replicate 2 |
| SAMN32973036 | F_DMS_10mM_rep1 | wt fluoride riboswitch multilength cotranscriptional DMS probing with 10 mM NaF, replicate 1 |
| SAMN32973037 | F_DMS_10mM_rep2 | wt fluoride riboswitch multilength cotranscriptional DMS probing with 10 mM NaF, replicate 2 |
| SAMN32973038 | G4P_BzCN_000uM_rep1 | wt ppGpp riboswitch multilength cotranscriptional BzCN probing with 0 uM ppGpp, replicate 1 |
| SAMN32973039 | G4P_BzCN_000uM_rep2 | wt ppGpp riboswitch multilength cotranscriptional BzCN probing with 0 uM ppGpp, replicate 2 |
| SAMN32973040 | G4P_BzCN_250uM_rep1 | wt ppGpp riboswitch multilength cotranscriptional BzCN probing with 250 uM ppGpp, replicate 1 |
| SAMN32973041 | G4P_BzCN_250uM_rep2 | wt ppGpp riboswitch multilength cotranscriptional BzCN probing with 250 uM ppGpp, replicate 2 |

**Supplementary Table 4 (continued)**
| SRA Acession | Sample Name | Experiment |
| --- | --- | --- |
| SAMN32973042 | G4P_DMS_000uM_rep1 | wt ppGpp riboswitch multilength cotranscriptional DMS probing with 0 uM ppGpp, replicate 1 |
| SAMN32973043 | G4P_DMS_000uM_rep2 | wt ppGpp riboswitch multilength cotranscriptional DMS probing with 0 uM ppGpp, replicate 2 |
| SAMN32973044 | G4P_DMS_250uM_rep1 | wt ppGpp riboswitch multilength cotranscriptional DMS probing with 250 uM ppGpp, replicate 1 |
| SAMN32973045 | G4P_DMS_250uM_rep2 | wt ppGpp riboswitch multilength cotranscriptional DMS probing with 250 uM ppGpp, replicate 2 |
| SAMN32973046 | G4P_EQ_BzCN_000uM_rep1 | wt ppGpp riboswitch multilength equilibrium BzCN probing with 0 uM ppGpp, replicate 1 |
| SAMN32973047 | G4P_EQ_BzCN_250uM_rep1 | wt ppGpp riboswitch multilength equilibrium BzCN probing with 250 uM ppGpp, replicate 1 |
| SAMN32973048 | G4PC69A_DMS_000uM_rep1 | ppGpp riboswitch C69A variant multilength cotranscriptional DMS probing with 0 uM ppGpp, replicate 1 |
| SAMN32973049 | G4PC69A_DMS_000uM_rep2 | ppGpp riboswitch C69A variant multilength cotranscriptional DMS probing with 0 uM ppGpp, replicate 2 |
| SAMN32973050 | G4PC69A_DMS_250uM_rep1 | ppGpp riboswitch C69A variant multilength cotranscriptional DMS probing with 250 uM ppGpp, replicate 1 |
| SAMN32973051 | G4PC69A_DMS_250uM_rep2 | ppGpp riboswitch C69A variant multilength cotranscriptional DMS probing with 250 uM ppGpp, replicate 2 |

**Supplementary Table 5.** RMDB data deposition table. Reactivity data generated in this work are freely available from the RNA Mapping Database (RMDB)^60^ (http://rmdb.stanford.edu), accessible using the RMDB ID numbers indicated in the table below.

| RMDB Accession | RNA | Experiment | Folding | Probe | Ligand | NTP Conc. | Concatenated | Smoothing |
| --- | --- | --- | --- | --- | --- | --- | --- | --- |
| CBAG4P_BZCN_0001 | wt ppGpp riboswitch | TECprobe-ML | Cotranscriptional | BzCN | none | 100uM | Reps 1 and 2 | Yes |
| CBAG4P_BZCN_0002 | wt ppGpp riboswitch | TECprobe-ML | Cotranscriptional | BzCN | 250uM ppGpp | 100uM | Reps 1 and 2 | Yes |
| CBAG4P_BZCN_0003 | wt ppGpp riboswitch | TECprobe-ML | Equilibrium | BzCN | none | 100uM | No | Yes |
| CBAG4P_BZCN_0004 | wt ppGpp riboswitch | TECprobe-ML | Equilibrium | BzCN | 250uM ppGpp | 100uM | No | Yes |
| CBAG4P_DMS_0001 | wt ppGpp riboswitch | TECprobe-ML | Cotranscriptional | DMS | none | 100uM | Reps 1 and 2 | Yes |
| CBAG4P_DMS_0002 | wt ppGpp riboswitch | TECprobe-ML | Cotranscriptional | DMS | 250uM ppGpp | 100uM | Reps 1 and 2 | Yes |
| CBAG4P_DMS_0003 | ppGpp riboswitch, C69A | TECprobe-ML | Cotranscriptional | DMS | none | 100uM | Reps 1 and 2 | Yes |
| CBAG4P_DMS_0004 | ppGpp riboswitch, C69A | TECprobe-ML | Cotranscriptional | DMS | 250uM ppGpp | 100uM | Reps 1 and 2 | Yes |
| CRCBFL_BZCN_0001 | wt fluoride riboswitch | TECprobe-ML | Cotranscriptional | BzCN | none | 100uM | Reps 1 and 2 | Yes |
| CRCBFL_BZCN_0002 | wt fluoride riboswitch | TECprobe-ML | Cotranscriptional | BzCN | 10mM NaF | 100uM | Reps 1 and 2 | Yes |
| CRCBFL_DMS_0001 | wt fluoride riboswitch | TECprobe-ML | Cotranscriptional | DMS | none | 100uM | Reps 1 and 2 | Yes |
| CRCBFL_DMS_0002 | wt fluoride riboswitch | TECprobe-ML | Cotranscriptional | DMS | 10mM NaF | 100uM | Reps 1 and 2 | Yes |
| PFLZTP_BZCN_0001 | wt ZTP riboswitch | TECprobe-ML | Cotranscriptional | BzCN | none | 100uM | Reps 1 and 2 | Yes |
| PFLZTP_BZCN_0002 | wt ZTP riboswitch | TECprobe-ML | Cotranscriptional | BzCN | 1mM ZMP | 100uM | Reps 1 and 2 | Yes |
| PFLZTP_DMS_0001 | wt ZTP riboswitch | TECprobe-ML | Cotranscriptional | DMS | none | 100uM | Reps 1 and 2 | Yes |
| PFLZTP_DMS_0002 | wt ZTP riboswitch | TECprobe-ML | Cotranscriptional | DMS | 1mM ZMP | 100uM | Reps 1 and 2 | Yes |
| PFLZTPS_BZCN_0001 | wt ZTP riboswitch | TECprobe-SL | Cotranscriptional | BzCN | none | 100uM | No | N/A |
| PFLZTPS_BZCN_0002 | wt ZTP riboswitch | TECprobe-SL | Cotranscriptional | BzCN | 1mM ZMP | 100uM | No | N/A |
| PFLZTPS_BZCN_0003 | wt ZTP riboswitch | TECprobe-SL | Cotranscriptional | BzCN | none | 100uM | No | N/A |
| PFLZTPS_BZCN_0004 | wt ZTP riboswitch | TECprobe-SL | Cotranscriptional | BzCN | 1mM ZMP | 100uM | No | N/A |
| PFLZTPS_BZCN_0005 | wt ZTP riboswitch | TECprobe-SL | Cotranscriptional | BzCN | none | 500uM | No | N/A |
| PFLZTPS_BZCN_0006 | wt ZTP riboswitch | TECprobe-SL | Cotranscriptional | BzCN | 1mM ZMP | 500uM | No | N/A |
| PFLZTPS_BZCN_0007 | wt ZTP riboswitch | TECprobe-SL | Cotranscriptional | BzCN | none | 500uM | No | N/A |
| PFLZTPS_BZCN_0008 | wt ZTP riboswitch | TECprobe-SL | Cotranscriptional | BzCN | 1mM ZMP | 500uM | No | N/A |

